# Oncogenic KRAS-MAPK signaling sustains a CD73 dependent therapeutic vulnerability in pancreatic cancer

**DOI:** 10.64898/2026.10.20.760559

**Authors:** Khadija Turabi, Urvinder Kaur Sardarni, Katherine E. Ankenbauer, Sabrina M. Torbit, Christopher P. Taranto, Shwetapadma Dash, Audrey Meade, Amelia T. Cephas, Kelsey A. Klute, Marina Pasca di Magliano, Thomas McFall, Kathleen E. DelGiorno, Jennifer M. Bailey-Lundberg

**Affiliations:** Department of Pathology, Microbiology, and Immunology, University of Nebraska Medical Center, Omaha, NE, USA; Fred & Pamela Buffett Cancer Center, University of Nebraska Medical Center, Omaha, NE, USA; Cancer Biology Program, Eppley Institute, University of Nebraska Medical Center, Omaha, NE, USA; Department of Cell and Developmental Biology, Vanderbilt University, Nashville, TN, USA; Program in Cancer Biology, Vanderbilt University, Nashville, TN, USA; Department of Internal Medicine, University of Nebraska Medical Center, Omaha, NE, USA; Department of Cell and Developmental Biology, University of Michigan, Ann Arbor, MI, USA; Department of Biochemistry, Medical College of Wisconsin and the LaBahn Pancreatic Cancer Program, Milwaukee, WI, USA; Cancer Center, Medical College of Wisconsin and the LaBahn Pancreatic Cancer Program, Milwaukee, WI, USA; Department of Surgery, Medical College of Wisconsin and the LaBahn Pancreatic Cancer Program, Milwaukee, WI, USA; Center for Computational Systems Biology, Vanderbilt University, Nashville TN, USA; Vanderbilt Ingram Cancer Center, Vanderbilt University Medical Center, Nashville, TN, USA; Vanderbilt Digestive Disease Research Center, Vanderbilt University Medical Center, Nashville, TN, USA

**Keywords:** Pancreatic ductal adenocarcinoma, KRAS, CD73, MAPK signaling, Tumor microenvironment, Tumor-associated macrophages, Desmoplasia, Combination therapy

## Abstract

Pancreatic ductal adenocarcinoma (PDAC) is predominantly driven by oncogenic KRAS signaling and sustained by a desmoplastic, immunosuppressive microenvironment limiting therapeutic response. CD73-mediated adenosine signaling supports this immune-excluded, tumor-supportive microenvironment. Whether dual targeting of KRAS-MAPK and CD73 signaling creates a therapeutic opportunity to improve tumor control remains unexplored. Analysis of *KRAS*-mutant cancer cell lines revealed increased expression of *NT5E* and adenosine receptor *ADORA2B*. Pharmacologic and genetic suppression of oncogenic KRAS inhibited CD73 expression through a MAPK-dependent program. In syngeneic KPC tumors, conversely, CD73 inhibition with quemliclustat increased ERK phosphorylation *in vivo*, suggesting adaptive MAPK activation following CD73 blockade. Combined CD73 and RAS(ON) inhibition *in vivo* produced the strongest antitumor response, reducing endpoint tumor volume beyond either monotherapy. This combination significantly suppressed tumor proliferation and αSMA-positive stroma and resulted in a substantial reduction in CD163-positive macrophage infiltration. RAS-MAPK targeted therapy was also associated with increased CD8α-positive T cell infiltration and granzyme B-positive staining. As therapeutic responses in PDAC have been linked to the gut microbiome, we further examined treatment-associated changes in the microbiome. RAS-MAPK inhibition produced the most pronounced microbial alterations, which correlated with tumor volume and immune markers. Together, these findings identify KRAS–MAPK driven CD73 signaling as a therapeutic vulnerability in PDAC and support combined CD73 and RAS(ON) inhibition as a strategy to suppress tumor growth, remodel stroma, and alleviate tumor-associated immune suppression.

## INTRODUCTION

Pancreatic ductal adenocarcinoma (PDAC) is one of the most therapeutically refractory malignancies. Activating mutations in *KRAS* occur in more than 90% of PDAC, making RAS signaling a dominant oncogenic dependency^1^. The recent development of inhibitors that target active, GTP-bound RAS has substantially changed the therapeutic landscape. Multi selective RAS(ON) inhibition has demonstrated robust antitumor activity in preclinical PDAC models^2,3^, while daraxonrasib has shown clinical activity in patients with previously treated RAS-mutant PDAC^4^. Recently, a randomized phase 3 trial showed significantly longer overall and progression free survival with daraxonrasib than with investigator’s choice of standard of care chemotherapy in previously treated metastatic PDAC patients^5^. Despite this progress, emerging resistance can be a major limitation. Analyses of tumors from patients treated with daraxonrasib have shown alterations in RAS, receptor tyrosine kinases, MAPK, and PI3K signaling, emphasizing the need for biologically informed combination strategies to enhance efficacy^6,7^.

The impact of oncogenic RAS signaling in PDAC extends beyond tumor cell proliferation and survival. Expression of mutant KRAS leads to transcriptional states and tumor derived signals that shape the surrounding microenvironment. Pharmacologic RAS inhibition can reorganize this microenvironment, and studies in immunocompetent models indicate that immune cells can substantially contribute to the extent of therapeutic response^8^. These observations infer that pro-tumorigenic signaling pathways downstream of RAS create therapeutic vulnerabilities between oncogenic signaling and the tumor microenvironment.

Extracellular adenosine signaling represents one such potential link between tumor cell signaling, metabolism, and immune regulation. CD39 hydrolyzes extracellular adenosine triphosphate (ATP) and adenosine diphosphate (ADP) to adenosine monophosphate (AMP), and the ecto-5’-nucleotidase CD73, encoded by the gene *NT5E*, converts AMP to adenosine. Adenosine can then signal through ADORA2A and ADORA2B adenosine receptors to suppress antitumor T cell and natural killer cell activity and to support immunoregulatory myeloid programs^9^. Our previous work in mouse models demonstrated that CD73– adenosine signaling differs according to the epithelial cell of origin of KRAS-driven PDAC^10^. Expression of oncogenic KRAS^G12V^ specifically in pancreatic ducts, as compared to acinar cells, results in increased *Nt5e* expression. As a result, duct-derived basal and squamous type PDAC exhibits higher CD73 expression and CD73 dependent adenosine mediated ADORA2B signaling which promotes both immune suppression and tumor progression, identifying a critical immune checkpoint in this molecular subtype of PDAC^10,11^. Clinically, the small molecule CD73 inhibitor quemliclustat was evaluated with chemotherapy and immune checkpoint blockade and was found to show efficacy. Tumor analyses showed modulation of adenosine-responsive transcriptional programs related to increased T cell activity^12^.

Despite these observations, how oncogenic RAS–MAPK and CD73 signaling interact in PDAC and whether this interaction represents a therapeutic vulnerability remains unclear. In this study, we investigated the relationship between oncogenic RAS-MAPK signaling and CD73 using transcriptomic analyses, inducible KRAS models, pathway-selective pharmacologic perturbation, and an immunocompetent PDAC model. Our findings identify a functional link between RAS–MAPK and CD73-associated signaling and provide a rationale for simultaneously targeting these pathways in PDAC.

## RESULTS

### Oncogenic KRAS drives a CD73 signaling program in PDAC

To identify targetable signaling pathways associated with oncogenic KRAS expression, we analyzed transcriptomic data from cancer cell lines available through cBioPortal^13–16^. Cell lines were stratified by *KRAS* mutation status and gene ontology and differentially expressed gene (DEG) analyses were performed (**Supplementary File 1**). Cell lines harboring mutations in *KRAS* showed enrichment of multiple nucleoside-related biological programs as compared with *KRAS* wild-type cell lines, including nucleoside metabolism, nucleoside biosynthesis, and nucleobase-containing processes (**Fig. 1A**). Among genes enriched in these pathways was *NT5E*, encoding CD73, which was found to be significantly elevated in *KRAS*-mutant cell lines regardless of point mutation (*KRAS^G12D^*, *KRAS^G12C^*, *KRAS^G12V^*) (**Fig. 1B**). The adenosine receptor *ADORA2B* was similarly increased (**Fig. 1C**), as was *FOSL1*, a known downstream transcriptional effector of RAS–MAPK signaling (**Supplementary Fig. 1A**). Together, these findings identify expression of multiple components of the CD73 pathway associated with mutant KRAS expression.

**Figure 1.**
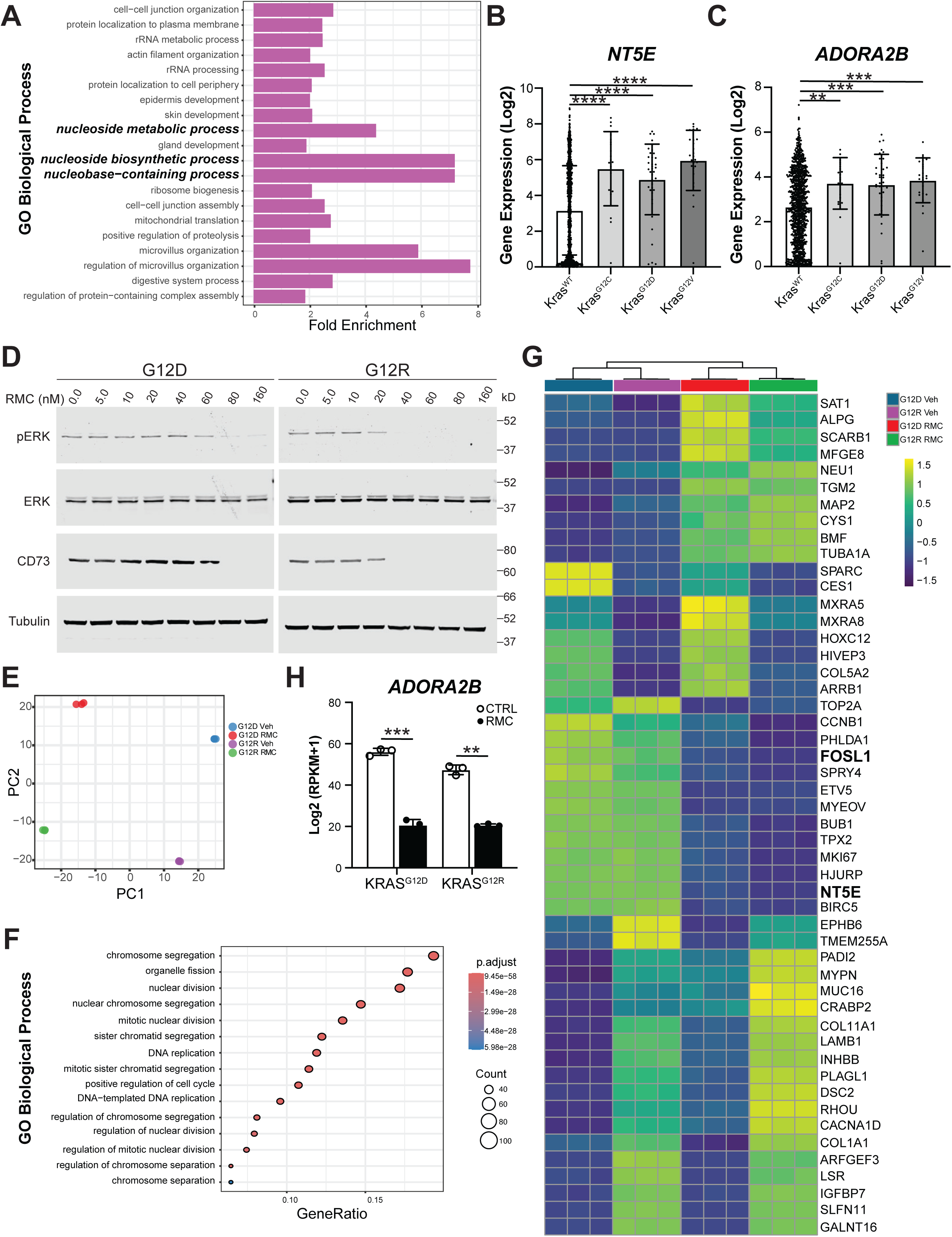
Oncogenic KRAS drives expression of adenosine signaling circuitry in cancer. (**A**) Gene ontology (GO) analysis of biological processes enriched in cBioPortal cancer cell line gene expression data in lines with wild-type *KRAS* (n = 889) vs oncogenic *KRAS* (G12D, n = 44; G12C, n = 21; G12V, n = 27) from multiple cancer types highlighting nucleoside biosynthesis and metabolism. Expression of (**B**) adenosine synthase *NT5E* and (**C**) adenosine receptor *ADORA2B* in the same lines. (**D**) Western blots of pancreatic cancer cell line PANC-1 bearing either a G12D or G12R point mutation treated with increasing concentrations of daraxonrasib for pERK1/2^Thr202+Tyr204^, total ERK, CD73, or alpha/beta-tubulin. (**E**) Principal component analysis (PCA) of RNA-seq of PANC-1 (G12D or G12R) cell lines treated with control or daraxonrasib. (**F**) GO analysis of RNA-seq in (E) showing a loss of proliferation-associated pathway gene expression in PANC-1 G12D cells with daraxonrasib treatment. (**G**) Heatmap with hierarchical clustering showing the top 50 changes in gene expression including a loss of *FOSL1* and *NT5E* with daraxonrasib. (**H**) Expression of adenosine receptor *ADORA2B* in the same treatment groups showing a loss of expression with daraxonrasib. Statistical significance was determined by a Welch’s -test for two-group comparisons or one-way ANOVA followed by Dunnett’s post-hoc test for multiple comparisons. **p < 0.01; ***p < 0.005; ****p < 0.001.

We next asked whether pharmacologic suppression of oncogenic KRAS signaling reverses this program. Treatment of syngeneic KRAS-mutant PANC-1 cell lines harboring either *KRAS^G12D^* or *KRAS^G12R^* with the RAS(ON) inhibitor daraxonrasib reduced ERK phosphorylation resulting in a concurrent decrease in CD73 protein expression with increasing drug concentration (**Fig. 1D**). In an orthogonal experiment, we treated both PANC-1 lines with either control or daraxonrasib and performed RNA-sequencing. Principal component analysis (PCA) demonstrated clear separation of experimental groups according to both *KRAS* genotype and treatment condition (**Fig. 1E**). Consistent with effective pathway inhibition, DEGs in treated lines were enriched for suppression of proliferative processes including chromosome segregation, nuclear division, DNA replication, and cell-cycle regulation (**Fig. 1F; Fig. S1F, Supplementary Files 2-3**). Daraxonrasib induced a shared transcriptional response across both KRAS-mutant models that included suppression of *FOSL1*, *NT5E*, and *ADORA2B* (**Fig. 1G-H; Fig. S1C-E).** Flow cytometric analysis further demonstrated a significant reduction in cell-surface CD73 expression following treatment with RAS(ON) inhibitors (**Fig. S1B**). Collectively, these findings demonstrate that oncogenic KRAS signaling drives CD73-ADORA2B pathway expression, which is effectively blocked with pharmacologic KRAS inhibition.

### Genetic loss of mutant KRAS diminishes *Nt5e*/CD73 expression

To determine whether maintenance of the CD73 program requires sustained oncogenic KRAS signaling, we examined two doxycycline-inducible *Kras^G12D^*murine pancreatic cancer cell lines (iKras#1 and iKras#2). Withdrawal of doxycycline, resulting in loss of oncogenic *Kras^G12D^* expression, significantly reduced known KRAS target gene *Fosl1* as well as *Nt5e* expression in both models (**Fig. 2A**). Consistent with these transcriptional changes, we identified loss of cell-surface CD73 protein abundance as measured by flow cytometry (**Fig. 2B**). These findings indicate that sustained oncogenic KRAS signaling is required to maintain *Nt5e*/CD73 expression.

**Figure 2.**
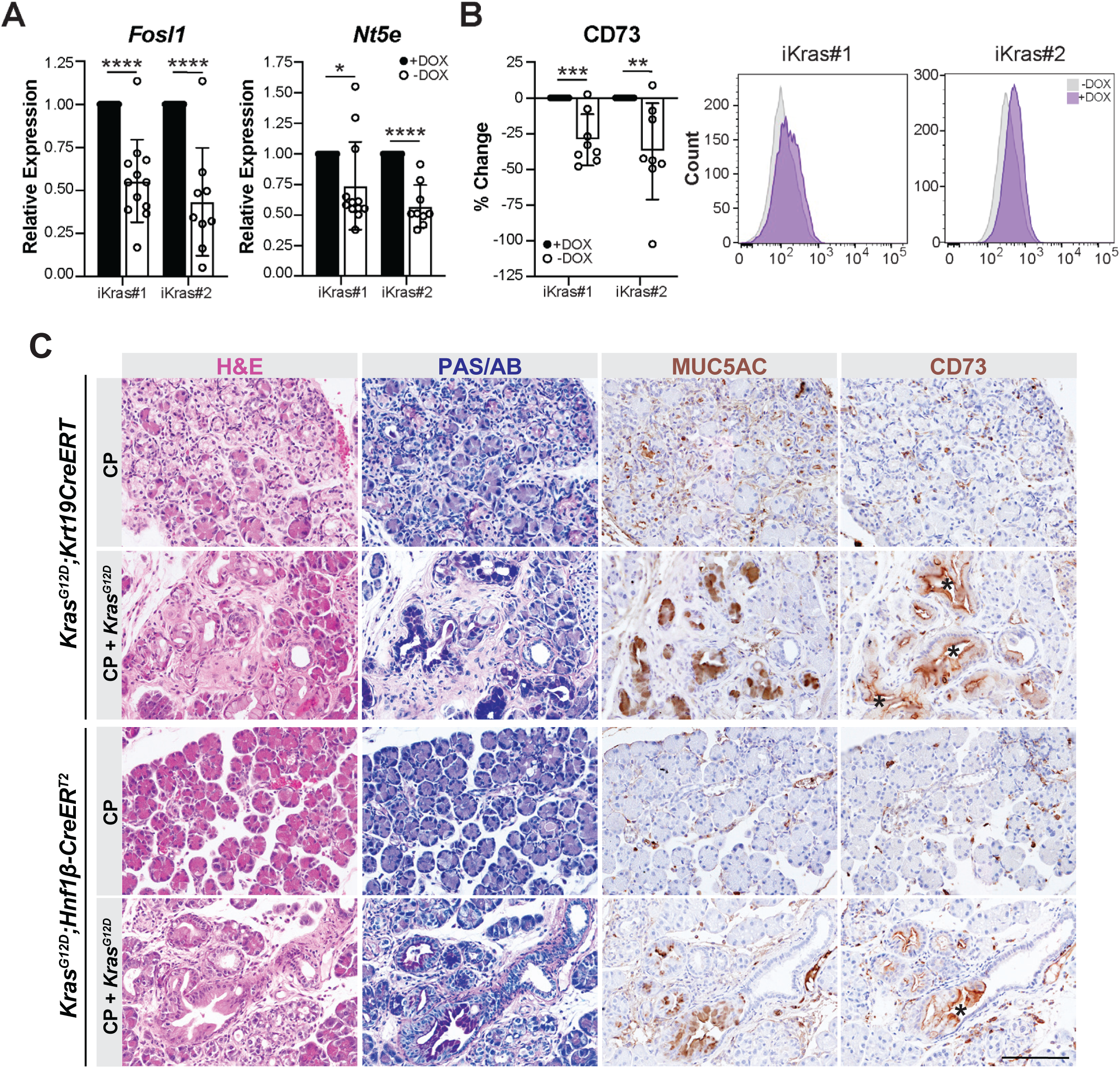
Oncogenic KRAS drives CD73 expression in murine models of pancreatic tumorigenesis and cancer. (**A**) Gene expression of known KRAS signaling target gene and effector *Fosl1* or *Nt5e* in two murine pancreatic cancer cell lines (iKras#1 and iKras#2) bearing a doxycycline-inducible *Kras^G12D^* allele. (**B**) CD73 protein expression in the same lines measured by flow cytometry. Statistical significance was determined by a Welch’s -test. (**C**) Histological analysis of murine pancreata from *Kras^G12D^* mice bred to ductal cell-specific inducible Cre alleles (either *Krt19CreERT* or *Hnf1*β*-CreER^T2^*) treated with caerulein to induce tissue injury and metaplasia or caerulein and tamoxifen to induce *Kras^G12D^*expression. H&E analysis demonstrating metaplasia in mice with caerulein alone (chronic pancreatitis, CP) and PanIN in mice with tamoxifen treatment, confirmed by mucin staining (Periodic Acid Schiff/Alcian Blue) and MUC5AC IHC. Enhanced CD73 expression is shown in PanIN by IHC. Scale bars are 50 μm. *p < 0.05; **p < 0.01; ***p < 0.005; ****p < 0.001.

We next investigated whether the KRAS–CD73 relationship was evident during pancreatic tumorigenesis *in vivo*. As we demonstrated in previous studies, caerulein-induced chronic pancreatitis resulted in metaplastic changes in the epithelium; subsequent activation of oncogenic KRAS by tamoxifen treatment promoted progression to pancreatic intraepithelial neoplasia (PanIN)^17^. PanIN lesions exhibited characteristic mucinous features, including PAS/Alcian blue and MUC5AC positivity, and demonstrated prominent epithelial CD73 expression (**Fig. 2C**). In contrast, CD73 expression was limited in the epithelium of chronic pancreatitis controls, lacking *Kras* activation. Together, these findings extend the association between oncogenic KRAS signaling and CD73 expression from cultured cells to early pancreatic neoplasia *in vivo*.

### Pharmacologic RAS–MAPK, but not PI3K, inhibition suppresses *NT5E*/CD73 expression

Having established a direct relationship between oncogenic KRAS and CD73 expression, we next asked what potentially targetable downstream signaling pathways drive CD73 expression. In the *KRAS^G12D^* mutant PANC-1 cell line, MRTX-1133 (*KRAS^G12D^* inhibitor) treatment significantly reduced expression of known target gene *FOSL1* (**Fig. 3A**) and *NT5E*/CD73 at both the gene and protein levels (**Fig.3A,D**) as well as adenosine receptor *ADORA2B* (**Fig. S2A-B**), whereas the *KRAS^G12C^* inhibitor AMG-510 had no effect, as expected. MEK inhibition with trametinib similarly suppressed gene and protein expression, while inhibition of PI3K signaling with wortmannin did not (**Fig. 3A,D; Fig. S2A-B**). In *KRAS^G12C^*mutant MIAPaCa-2 cells, MRTX-133 had no effect while AMG-510 (KRAS^G12C^ inhibitor) significantly reduced *FOSL1*, *NT5E*/CD73, and *ADORA2B*; treatment with trametinib yielded similar results, while wortmannin treatment had no effect (**Fig. 3B,E; Fig. S2A-B**). In BxPC3 cells, which carry a BRAF in-frame deletion that constitutively activates MAPK (and wild-type *KRAS*), neither *KRAS* directed inhibitor suppressed *FOSL1* or *NT5E*/CD73, whereas trametinib markedly reduced gene and protein expression (**Fig. 3C,F; Fig. S2A-B**). In contrast to PANC-1 and MIAPaCa-2 cell lines, loss of *ADORA2B* expression was observed in BxPC3 cells in response to both trametinib and wortmannin treatment (**Fig. S2A-B**). Immunoblotting confirmed modulation of the intended signaling pathways following treatment, including suppression of ERK phosphorylation by MAPK-directed therapies and inhibition of AKT phosphorylation by wortmannin (**Fig. S2C-F**). Overall, loss of *NT5E/*CD73 expression by mutant KRAS targeted therapeutics is largely accomplished by suppressing MAPK, but not PI3K, signaling.

**Figure 3.**
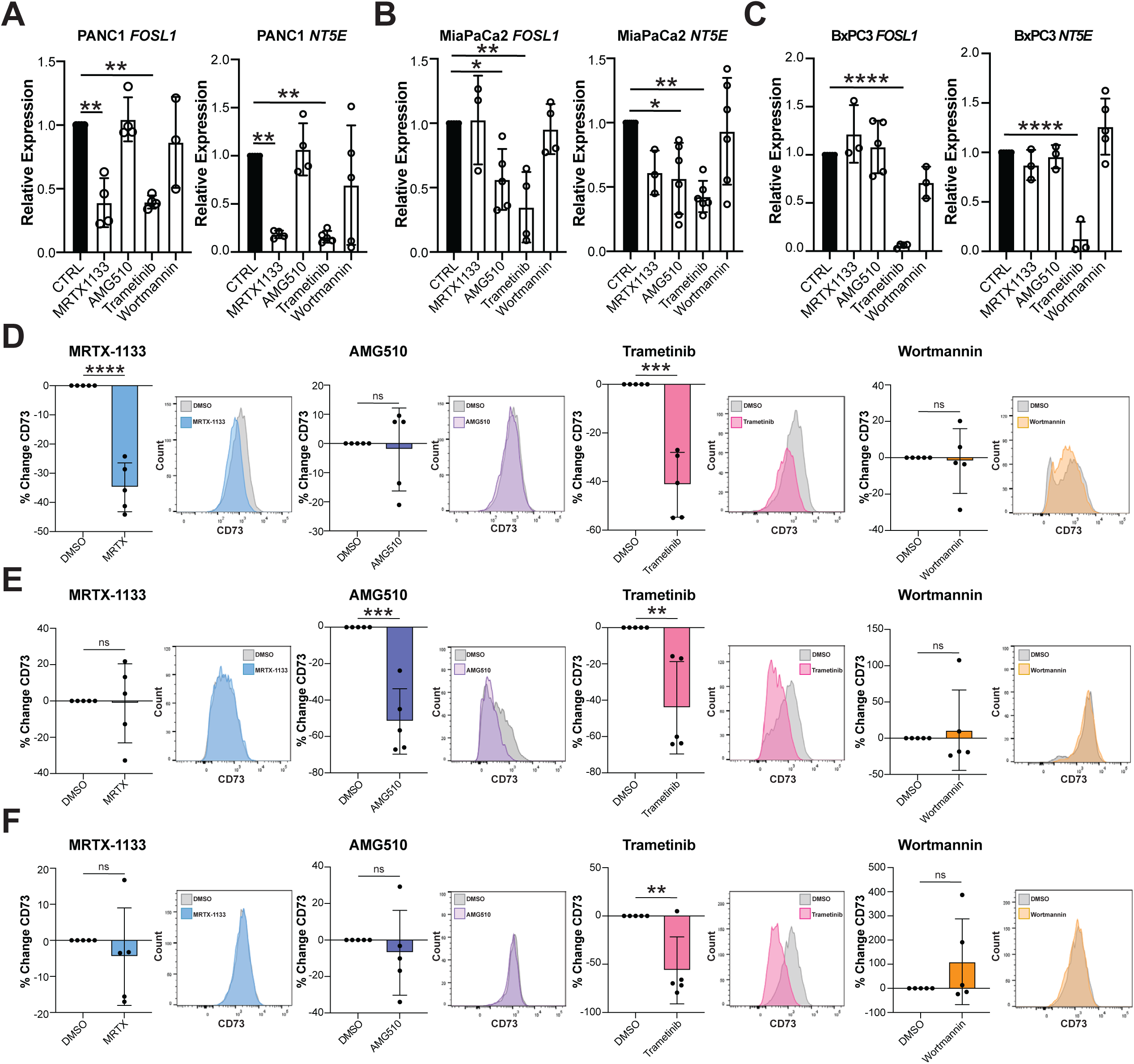
Oncogenic KRAS drives CD73 expression through MAPK, but not PI3K signaling. Expression of mutant KRAS signaling downstream target gene *FOSL1* or *NT5E* as measured by qRT-PCR in (**A**) PANC-1 (KRAS^G12D^), (**B**) MIAPaCa-2 (KRAS^G12C^), or (**C**) BxPC3 (KRAS wild-type) pancreatic cancer cell lines treated with control (CTRL) or KRAS^G12D^ inhibitor MRTX-1133, KRAS^G12C^ inhibitor AMG-510, MEK inhibitor trametinib, or PI3K inhibitor wortmannin for 48 h. CD73 protein expression in (**D**) PANC-1, (**E**) MIAPaCa-2, or (**F**) BxPC3 cell lines treated with the same drugs as measured by flow cytometry. Statistical significance was determined by a Welch’s -test for two-group comparisons or one-way ANOVA followed by Dunnett’s post-hoc test for multiple comparisons. ns, not significant; *p < 0.05; **p < 0.01; ***p < 0.005; ****p < 0.001.

### Efficacy of combined CD73 and RAS–MAPK inhibition *in vivo*

Having shown through transcriptomic, genetic, and pharmacologic approaches that suppression of oncogenic KRAS-MAPK signaling reduces CD73 expression, including *FOSL1* and *NT5E*/CD73 expression, we next asked whether simultaneous targeting of these pathways could improve therapeutic efficacy *in vivo*. PDAC tumors were generated subcutaneously using cell lines derived from *Pdx1-Cre;LSL-Kras^G12D/+^;LSL-Trp53^R^*^172^*^H/+^* (KPC) mice, and were subsequently treated with vehicle, the CD73 inhibitor quemliclustat, the RAS(ON) inhibitor daraxonrasib, the MEK inhibitor trametinib, or combinations of quemliclustat with daraxonrasib (Q + D) or trametinib (Q + T) (**Fig. 4A**). Quemliclustat monotherapy produced a modest reduction in endpoint tumor volume compared with vehicle, whereas inhibition of KRAS/MAPK signaling with either daraxonrasib or trametinib markedly suppressed tumor growth. Notably, Q + D produced the greatest reduction in tumor volume and further decreased tumor burden relative to daraxonrasib monotherapy, whereas the combination of quemliclustat with trametinib did not significantly reduce tumor volume compared to the trametinib monotherapy (**Fig. 4B**). Tumor histology was evaluated by hematoxylin and eosin (H&E) staining (**Fig. 4C,D**).

**Figure 4.**
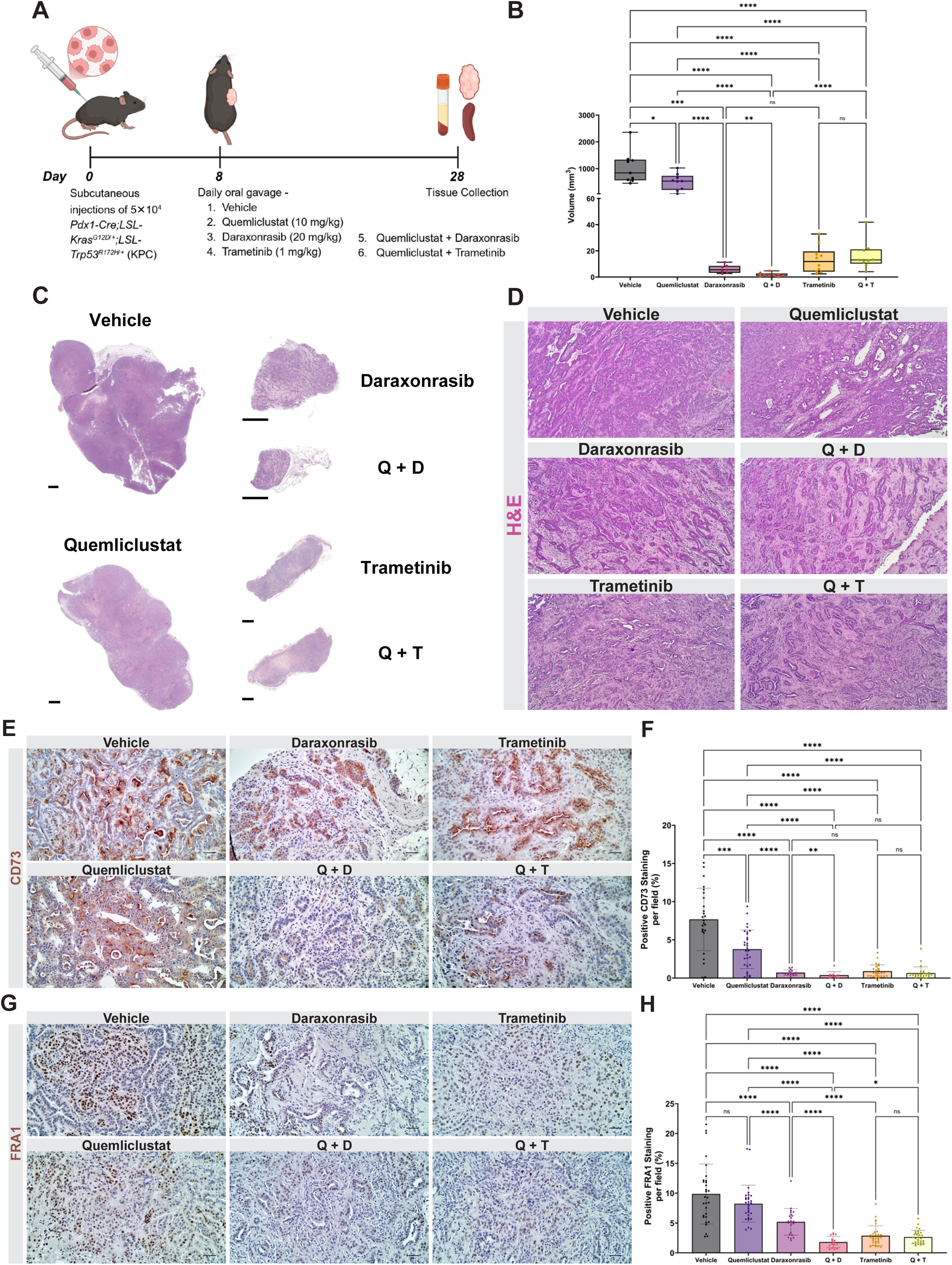
CD73 signaling inhibition potentiates RAS(ON) inhibition to suppress KPC pancreatic tumor growth *in vivo*. (**A**) Schematic of the subcutaneous KPC pancreatic tumor model and treatment regimen. Mice were inoculated subcutaneously with 5 × 10^4^ KPC cells and treatment was initiated on day 8 with vehicle, quemliclustat (10 mg/kg), daraxonrasib (20 mg/kg), trametinib (1 mg/kg), quemliclustat plus daraxonrasib (Q + D), or quemliclustat plus trametinib (Q + T). Tumors were harvested on day 28. (**B**) Box and whisker plot (median, first and third quartiles, min and maximum) showing endpoint tumor volumes for vehicle (n=10), quemliclustat (n=9), daraxonrasib (n=7), Q + D (n=9), trametinib (n=10), and Q + T (n=9). (**C**) Representative whole-tumor H&E-stained scanned sections from each treatment group. Scale bars are 1 mm. (**D**) Representative H&E images demonstrating tumor histopathology across treatment groups. Scale bars are 50 μm. (**E**) Representative images of CD73 IHC staining of mouse subcutaneous tumor sections from the different treatment groups. Scale bars are 50 μm. (**F**) Quantification of the percentage positive CD73 staining per field in each group. (**G**) Representative images of FRA1 IHC staining of mouse subcutaneous tumor sections from the different treatment groups. Scale bars are 50 μm. (**H**) Quantification of the percentage positive FRA1 staining per field in each group. All quantifications were performed for n = 3 mice per treatment group, and 5–10 fields per mouse were analyzed. Statistical significance was determined by Mann–Whitney U test. ns, not significant; *p < 0.05; **p *<* 0.01; ***p < 0.001; ****p < 0.0001.

We next determined, using immunohistochemistry (IHC), whether the KRAS–CD73 relationship identified *in vitro* was sustained *in vivo* in drug-treated tumors. There was a significant reduction in CD73 expression following all monotherapies. Quemliclustat in combination with daraxonrasib led to a further decline in CD73 expression; however, the combination with trametinib did not significantly reduce expression in comparison with trametinib monotherapy (**Fig. 4E,F**). As these therapies resulted in marked reduction in CD73 expression within the tumors, we next examined upstream expression of CD39. CD39 levels, however, remained unchanged across all treatment groups (**Fig. S3A,B**). We also evaluated vascular endothelial staining using CD31. CD31-positive area was significantly reduced following trametinib monotherapy and combination treatment with quemliclustat plus either daraxonrasib or trametinib, while neither quemliclustat nor daraxonrasib monotherapy significantly altered CD31-positive area (**Fig. S3C,D**). Furthermore, consistent with *in vitro* results, the KRAS effector FRA1 (encoded by the gene *FOSL1*) was reduced by direct KRAS–MAPK pathway inhibition, with the lowest levels observed in the daraxonrasib combination treatment group. In contrast, like the control tumors, FRA1 expression was sustained following quemliclustat monotherapy (**Fig. 4G,H**). Together, these findings extend the KRAS–MAPK-CD73/FRA1 relationship observed *in vitro* to an *in vivo* tumor context and demonstrate robust antitumor activity following simultaneous targeting of these pathways.

### Adaptive MAPK signaling and suppression of proliferative and stromal programs following CD73 inhibition

Given the reciprocal relationship between CD73 and KRAS–MAPK signaling observed in our *in vitro* studies, we next evaluated MAPK pathway activity following therapeutic CD73 inhibition. Quemliclustat monotherapy significantly increased pERK1/2 staining relative to vehicle, consistent with an adaptive response to CD73 blockade (**Fig. 5A,B**). In contrast, pERK1/2 levels were reduced when quemliclustat was combined with trametinib, consistent with MEK inhibition suppressing this treatment-associated MAPK activation. However, there was no significant change in pERK1/2 expression with daraxonrasib treatment (**Fig. 5A,B**).

**Figure 5.**
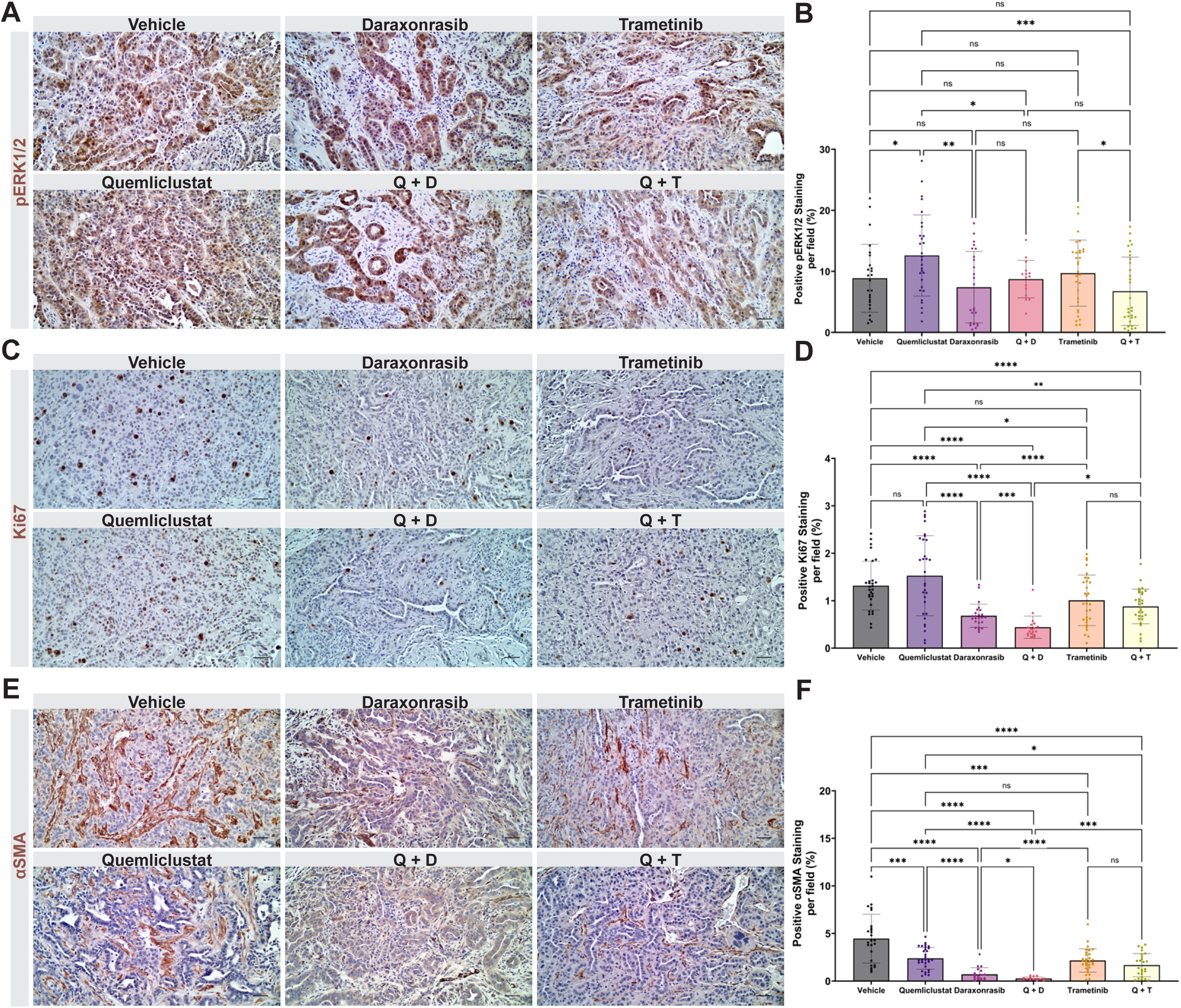
CD73 and KRAS–MAPK signaling inhibition drives MAPK pathway adaptation with reduced tumor proliferation and stromal activation. Representative images of (A) pERK1/2^Thr202/Thr185^, (C) Ki67 and (**E**) αSMA IHC staining of mouse subcutaneous tumor sections from the different treatment groups. Scale bars are 50 μm. Quantification of the percentage positive of (**B**) pERK1/2, (**D**) Ki67 and (**F**) αSMA staining per field in each group. All quantifications were performed for n = 3 mice per treatment group, and 5–10 fields per mouse were analyzed. Statistical significance was determined by Mann–Whitney U test. ns, not significant, *p < 0.05; **p < 0.01; ***p < 0.001; ****p < 0.0001.

We subsequently examined whether these signaling pathway changes were associated with altered tumor cell proliferation. Despite its effect on overall tumor burden and CD73 expression, quemliclustat alone did not significantly reduce Ki67 staining relative to vehicle. In contrast, daraxonrasib and trametinib significantly decreased tumor cell proliferation, with the most pronounced reduction observed following Q + D treatment (**Fig. 5C,D**). These findings are consistent with the marked tumor growth inhibition observed *in vivo* and indicate that suppression of KRAS–MAPK signaling, in comparison to CD73 blockade alone, robustly inhibit tumor cell proliferation.

As we’ve shown that CD73 derived adenosine can also influence the tumor stroma^10^, we assessed αSMA-positive fibroblast content across treatment groups. Quemliclustat reduced αSMA staining compared with vehicle, while substantially greater reductions were observed following KRAS–MAPK pathway inhibition. Strikingly, αSMA-positive staining was nearly abrogated in Q + D treated tumors and was also reduced following Q + T treatment (**Fig. 5E,F**). Collectively, these findings demonstrate that CD73 inhibition is accompanied by adaptive MAPK activation *in vivo*, whereas concurrent targeting of KRAS–MAPK signaling produces broader suppression of tumor proliferative and stromal programs.

### Cytotoxic immune remodeling and marked reduction of CD163-positive macrophages following CD73 and KRAS targeting

We next investigated whether the antitumor effects of CD73 and KRAS–MAPK pathway inhibition were accompanied by remodeling of the tumor immune microenvironment. CD8α-positive T cell infiltration was increased across treatment groups relative to vehicle, with the greatest accumulation observed in tumors treated with trametinib or Q + T (**Fig. 6A,B**). The combination of quemliclustat with either daraxonrasib or trametinib, however, did not substantially increase CD8α-positive T cells compared to the daraxonrasib or trametinib monotherapy. Granzyme B staining, commonly associated with cytotoxic lymphocyte activity, was also increased in several RAS–MAPK targeted groups, with a particularly prominent increase following daraxonrasib treatment (**Fig. 6C,D**).

**Figure 6.**
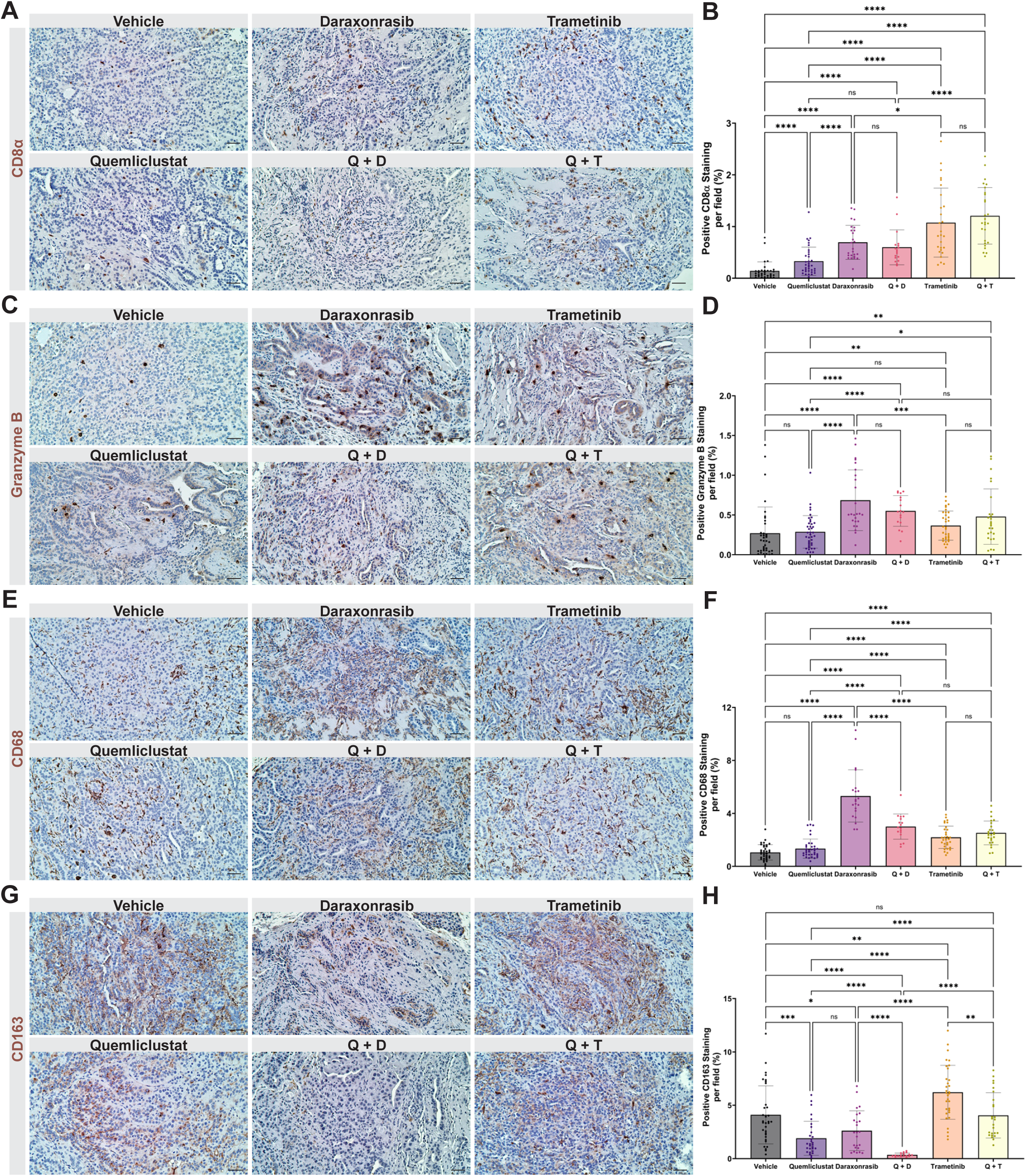
Enhanced cytotoxic immune infiltration and pronounced reduction of CD163⁺ macrophages with the dual targeting strategy. Representative images of (**A**) CD8α, (**C**) Granzyme B, (**E**) CD68, and (**G**) CD163 IHC staining of mouse subcutaneous tumor sections from the different treatment groups. Scale bars are 50 μm. Quantification of the percentage positive of (**B**) CD8α, (**D**) Granzyme B, (**F**) CD68, and (**H**) CD163 staining per field in each group. All quantifications were performed for n = 3 mice per treatment group, and 5– 10 fields per mouse were analyzed. Statistical significance was determined by Mann–Whitney U test. ns, not significant, *p < 0.05; **p < 0.01; ***p < 0.001; ****p < 0.0001.

The various therapies also produced marked changes in macrophage infiltration. Total CD68-positive macrophage staining was substantially increased following daraxonrasib treatment and remained elevated with Q + D, with more moderate increases observed following trametinib based treatment (**Fig. 6E,F**). In contrast, CD163-positive immunosuppressive macrophages exhibited a distinct treatment dependent pattern. Quemliclustat reduced CD163 staining relative to vehicle, whereas Q + D produced a striking near-complete loss of CD163 expression, representing the most pronounced reduction among all treatment groups (**Fig. 6G,H**). Conversely, trametinib monotherapy was associated with increased CD163 staining, an effect that was significantly attenuated by the addition of quemliclustat. Thus, although KRAS–MAPK inhibition increased overall macrophage abundance, concurrent CD73 blockade reduced immunosuppressive macrophages, most prominently following quemliclustat and daraxonrasib combined treatment.

These *in vivo* findings reveal a symbiotic therapeutic interaction among these pathways. CD73 blockade induces adaptive MAPK signaling, while combining CD73 inhibition with either RAS(ON) or MEK inhibition enhances tumor control and is accompanied by suppression of tumor proliferation and αSMA-positive stroma, increased cytotoxic immune infiltration, and decreased abundance of immunosuppressive macrophages.

### Treatment-associated alterations in gut microbial composition following CD73 and KRAS–MAPK pathway inhibition

Previous studies in PDAC have shown that treatment can alter the gut microbiome^18,19^, while microbial composition and microbiota-derived metabolites can influence therapeutic responses^20,21^. Particularly, CD73-dependent adenosine production has been implicated in intestinal host–microbiome interactions, while MEK inhibition has been shown to alter gut microbial composition in preclinical studies^22,23^. We therefore examined whether CD73 and KRAS–MAPK inhibition altered gut microbial composition and whether these changes were associated with tumor volume and immune markers. To address this, we collected stool from treated mice at endpoint and performed 16S rRNA gene amplicon sequencing. Analysis of alpha diversity demonstrated treatment-specific effects. Microbial richness, measured by observed amplicon sequence variants (ASVs), was reduced following quemliclustat treatment, whereas trametinib and Q + T treatment increased richness relative to vehicle controls (**Fig. 7A**). Shannon diversity was similarly increased following trametinib treatment (**Fig. 7B**).

**Figure 7.**
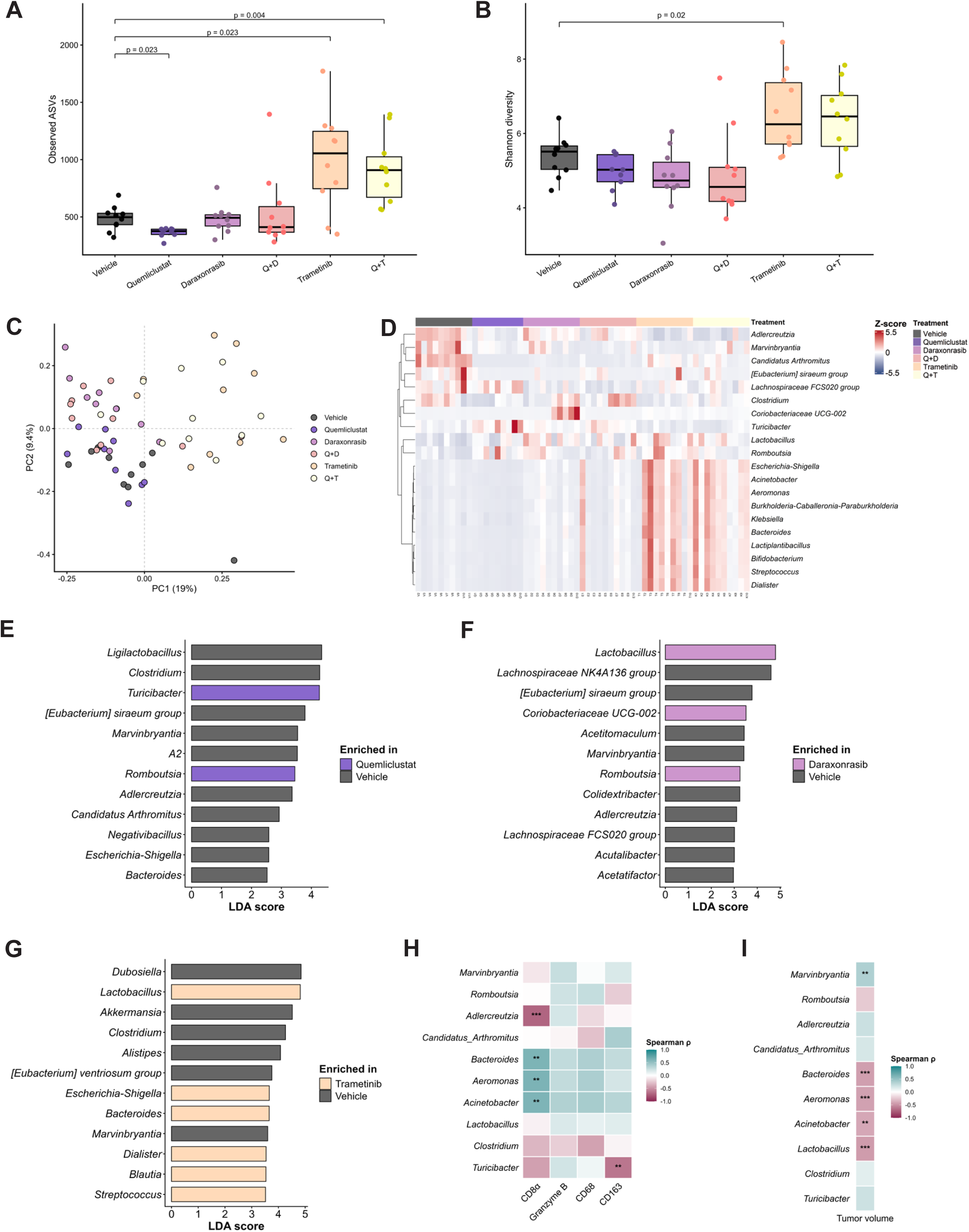
CD73 and KRAS–MAPK pathway inhibition are associated with changes in the gut microbiome. Alpha diversity measured by (**A**) observed ASVs and (**B**) Shannon diversity across vehicle, quemliclustat, daraxonrasib, quemliclustat + daraxonrasib (Q + D), trametinib and quemliclustat + trametinib (Q + T) treated mice. Statistical significance was determined using pairwise Wilcoxon rank-sum tests with Benjamini–Hochberg adjustment for multiple comparisons. (**C**) Principal coordinate analysis (PCoA) based on Bray–Curtis distances showing differences in microbial community composition across treatment groups. Statistical significance was determined using PERMANOVA and group dispersion was evaluated using PERMDISP. (**D**) Heat map showing row-wise Z-scores of relative abundance for the 20 genera that were differentially abundant when each monotherapy group was compared with the vehicle group. Columns represent individual mice grouped by treatment. LEfSe analysis showing the 12 genera with the highest absolute LDA scores for (**E**) vehicle versus quemliclustat, (**F**) vehicle versus daraxonrasib and (**G**) vehicle versus trametinib. Bar color indicates the group in which each genus was enriched. LEfSe significance was defined as a Kruskal–Wallis p < 0.05 and an absolute LDA score ≥ 2. Spearman correlation heat map showing associations between the top 10 selected bacterial genera and (**H**) CD8α, Granzyme B, CD68, CD163 and (**I**) tumor volume. Correlations with immune markers were performed for mice that had both microbiome and IHC data (n = 3 per treatment group), whereas correlations with tumor volume were performed using all mice (n = 59). Color represents the Spearman correlation coefficient (ρ). *p < 0.05; **p < 0.01; ***p < 0.001.

We next assessed differences in overall community structure using Bray-Curtis dissimilarity. Principal coordinate analysis demonstrated significant treatment-associated differences in microbial composition (PERMANOVA *p* = 0.001; PERMDISP *p* = 0.28) (**Fig. 7C**). Differential abundance analyses further revealed distinct microbial signatures across treatment groups (**Fig. 7D-G**). Quemliclustat treatment was associated with enrichment of genera including *Turicibacter* and *Romboutsia,* whereas daraxonrasib treatment enriched *Lactobacillus, Coriobacteriaceae UCG-002*, and *Romboutsia.* Trametinib produced the broadest taxonomic alterations and was associated with enrichment of multiple genera, including *Lactobacillus, Bacteroides, Dialister, Blautia,* and *Streptococcus* (**Fig. 7E-G**). Notably, addition of quemliclustat to either daraxonrasib or trametinib did not identify additional differentially abundant genera relative to the corresponding KRAS–MAPK inhibitor monotherapies. Together, these findings indicate that CD73- and KRAS–MAPK-targeted therapies are associated with distinct alterations in microbial diversity and community composition, with the most extensive changes observed following trametinib treatment.

To explore associations between treatment-associated microbial changes and the tumor microenvironment, we performed Spearman correlation analyses between differentially abundant genera and immune and tumor phenotypes. Bacteroides, Aeromonas, and Acinetobacter positively correlated with CD8α-positive T cell infiltration, whereas Adlercreutzia was negatively associated with CD8α (**Fig. 7H**). Turicibacter was negatively associated with CD163-positive macrophages, while Bacteroides, Aeromonas, Acinetobacter, and Lactobacillus were negatively correlated with tumor volume and Marvinbryantia was positively associated (**Fig. 7H**). Together, these exploratory findings demonstrate that CD73- and KRAS–MAPK-targeted therapies are accompanied by alterations in gut microbial composition and identify associations between specific microbial taxa, tumor burden, and immune features of the tumor microenvironment. Whether these microbial alterations contribute to therapeutic response or arise as a consequence of treatment and tumor control remains to be determined.

## DISCUSSION

Previous studies from our group demonstrated that CD73 promotes adenosine accumulation in pancreatic cancer^10^. The current study extends those observations by identifying oncogenic KRAS as an upstream regulator of a CD73-associated transcriptional program and by demonstrating therapeutic benefit from targeting both pathways simultaneously. Specifically, oncogenic KRAS–MAPK signaling supports expression of *NT5E*/CD73, *FOSL1*, and *ADORA2B*, whereas pharmacologic CD73 inhibition is accompanied by increased ERK phosphorylation *in vivo*. Higher CD73 expression in tumors that respond poorly to KRAS^G12D^ inhibition further supports a relationship between these pathways in the context of therapeutic response. These findings suggest that KRAS–MAPK and CD73 signaling are functionally linked in PDAC and reveal a therapeutic vulnerability that can be exploited through combined pathway inhibition.

CD73 is an established driver of pancreatic tumor progression and immune suppression, promoting extracellular adenosine production and shaping a tumor-supportive microenvironment^10,24^. However, the relationship between CD73 and the oncogenic signaling programs that sustain PDAC is poorly understood. Across transcriptomic, genetic, and pharmacologic models, we found that suppression of KRAS–MAPK signaling consistently reduced expression of CD73-associated genes, including *NT5E* and *ADORA2B*, while also reducing cell-surface CD73 abundance. Together, these findings suggest that CD73 forms part of a broader KRAS–MAPK-dependent transcriptional program and raise the possibility that its regulation is mediated through downstream ERK-responsive transcription factors.

The accompanying changes in *FOSL1* are particularly interesting given the established role of AP-1 factors, including FRA1 encoded by *FOSL1*, in KRAS-driven transcriptional and enhancer programs during pancreatic tumorigenesis^25,26^. Suppression of KRAS–MAPK signaling consistently reduced *FOSL1* expression in parallel with *NT5E*, raising the possibility that components of the AP-1 transcriptional network contribute to regulation of the CD73 program. This is consistent with prior evidence in melanoma that MAPK-dependent c-Jun/AP-1 activity directly promotes NT5E transcription through binding to an intronic enhancer^27^. Although the present study does not establish a direct mechanistic role for FOSL1, these findings provide a framework for future studies aimed at defining how oncogenic KRAS controls CD73 expression and related signaling pathways.

One of the most notable findings of this study was the increase in ERK phosphorylation following CD73 inhibition.

Adaptive reactivation of RAS–MAPK signaling is a well-recognized mechanism of resistance to targeted therapies, including RAF and KRAS^G12C^ inhibition, often through relief of pathway feedback and upstream RTK–RAS signaling^6^. More recently, allele-specific mechanisms that restore or maintain RAS pathway signaling have been identified in daraxonrasib-resistant PDAC^7^. Although CD73 blockade acts outside this canonical signaling cascade, the increased ERK phosphorylation observed here raises the possibility of a compensatory response whose mechanism and contribution to therapeutic efficacy require further investigation. The superior tumor control achieved by this combination is particularly relevant given the ongoing clinical development of both agents in PDAC^4,5,12^. Interestingly, addition of quemliclustat to trametinib did not further improve tumor control, suggesting that the consequences of CD73 co-targeting may depend on where the pathway is inhibited within the RAS–MAPK signaling cascade. Understanding the basis for these differences may provide further insight into mechanisms of therapeutic adaptation and combination response.

Combined pathway inhibition also produced substantial changes within the tumor microenvironment. The marked reduction in αSMA-positive stroma suggests that simultaneous targeting of KRAS–MAPK and CD73 signaling influences fibroblast-associated programs, although αSMA alone cannot distinguish changes in fibroblast abundance from changes in fibroblast state^28^. Consistent with previous reports, RAS–MAPK inhibition was additionally associated with increased CD8α and granzyme B staining, supporting enhanced immune engagement within the tumor microenvironment^8^. Recent studies of MRTX1133 and daraxonrasib, however, have shown that KRAS inhibition can induce a spectrum of effector, exhausted, and regulatory T cell states rather than a uniformly productive antitumor response^29^. These observations suggest that although KRAS inhibition promotes immune infiltration, suppressive mechanisms are likely to persist. Recent work further demonstrated that CDK8-dependent remodeling of the tumor microenvironment promotes resistance to KRAS^G12D^ inhibition through increased CXCL2 secretion and reduced FAS expression^30^. These findings emphasize that the initial immune effects of KRAS inhibition may be altered as tumors adapt to treatment. Given the established immunoregulatory effects of extracellular adenosine^9,10^, CD73 signaling may represent one pathway contributing to this residual immune suppression and therefore an attractive target for combination therapy.

Changes in macrophage infiltration with KRAS–MAPK and CD73 targeting were particularly striking. Daraxonrasib increased overall CD68-positive macrophage abundance, while quemliclustat alone had little effect and attenuated this increase in combination. This pattern is consistent with emerging evidence that KRAS-targeted therapies substantially reshape the myeloid compartment in PDAC. Pharmacologic KRAS inhibition has been associated with increased macrophage infiltration together with reduced immunosuppressive macrophage states^31^, while genetic studies suggest that sustained oncogenic KRAS signaling promotes pro-tumorigenic macrophage programs^32^. Notably, combined CD73 and RAS inhibition produced a pronounced reduction in CD163-positive macrophages, a population commonly associated with immunoregulatory tumor-associated macrophage states. This raises the possibility that CD73-associated signaling contributes to the macrophage response that emerges during KRAS-targeted therapy. Whether this reflects a direct effect of adenosine signaling on macrophages or arises through secondary changes in the tumor microenvironment remains an important area for future investigation.

In addition to remodeling the tumor microenvironment, CD73 and KRAS–MAPK pathway inhibition were associated with changes in the gut microbiome. Similar treatment-associated microbial shifts have been reported in PDAC^18,19^. More recently, MEK inhibition was shown to alter gut microbial composition and microbial arginine metabolism in colorectal cancer models, suggesting their role in antitumor immunity^23^. Although KRAS inhibition has been shown to remodel the immune microenvironment in PDAC^29^, its association with gut microbial composition remains poorly characterized. In our study, microbial changes were most pronounced following KRAS–MAPK inhibition. Furthermore, *Lactobacillus* was enriched following RAS– MAPK directed treatment and was negatively associated with tumor volume. This observation is consistent with a recent study in which higher fecal *Lactobacillus* abundance was associated with improved progression-free and overall survival in patients with PDAC, while *Lactobacillus reuteri* reduced tumor growth in preclinical models and increased intratumoral NK cell infiltration^33^. Other *Lactobacillus* based therapies have also been reported to enhance antitumor immune responses in pancreatic cancer models^34^. Beyond changes in microbial composition, microbiota-derived metabolites may also play a role in antitumor immunity. Notably, bacterial inosine has been shown to enhance responses to immune checkpoint blockade through adenosine A2A receptor-dependent T cell, suggesting that microbial purine metabolism may contribute to immune regulation^35^. Although the present data do not establish a causal contribution of microbial alterations to therapeutic efficacy, correlations between specific bacterial genera, tumor burden, CD8α-positive lymphocytes, and CD163-positive macrophages suggest that treatment-induced microbial remodeling may be associated with changes in the tumor microenvironment.

Interestingly, microbial changes appeared to be driven largely by RAS–MAPK inhibition, as adding quemliclustat to either daraxonrasib or trametinib produced few additional changes in microbial composition compared with the corresponding monotherapy. This contrasted with the tumor microenvironment, where CD73 blockade had more apparent effects when combined with RAS–MAPK inhibition, particularly within the stromal and CD163-positive macrophage compartments. Thus, the systemic microbial response and intratumoral response to these therapies may not occur in parallel, suggesting that CD73 inhibition exerts its strongest additional effects locally within the tumor rather than through further remodeling of the gut microbiome.

Several limitations should be acknowledged. Although prior studies from our group demonstrated that CD73 contributes to adenosine accumulation in PDAC, extracellular adenosine concentrations were not directly quantified in the current study. Additionally, the mechanisms responsible for increased MAPK activity following CD73 inhibition remain undefined. As these inhibitors could have direct effects on the immune and stromal cells, this limits our ability to attribute the observed microenvironmental changes to tumor cell intrinsic KRAS– CD73 signaling alone. Furthermore, the microbiome findings are correlative and do not establish whether treatment-associated microbial changes contribute to therapeutic response or arise secondarily from changes in tumor burden or host physiology. Functional studies will be needed to determine whether the identified microbial populations influence treatment efficacy. Future studies will determine how changes in adenosine signaling through ADORA2B influence stromal and immune remodeling and whether these effects contribute directly to adaptive MAPK activation. Testing combined CD73 and RAS inhibition in orthotopic and genetically engineered models of established PDAC will further clarify the therapeutic relevance of this relationship.

Collectively, our findings support a model in which oncogenic KRAS–MAPK signaling sustains a CD73-associated transcriptional program, while CD73 inhibition is accompanied by increased MAPK activity. This relationship creates a therapeutic vulnerability that can be exploited through combined pathway inhibition. As KRAS-targeted therapies become increasingly incorporated into the treatment landscape of PDAC, co-targeting CD73 may represent a strategy to enhance tumor control while simultaneously remodeling stromal and immune components of the tumor microenvironment.

## METHODS

### cBioPortal Analysis

mRNA expression (log2) values and mutation data were downloaded from cBioPortal (ccle_broad_2019) for cancer cell lines^13–16^. Data analyses were executed using R (version 4.4.2, 2024-10-31 ucrt), tailored for a 64-bit windows platform. R integrated package utilized in the study consisted of ‘limma(‘3.62.2’)’, edgeR(’34.4.2’)’, and clusterProfiler(‘4.14.6’). Samples were classified as *KRAS*-mutant or wild-type based on the provided mutation annotations. Expression matrices were imported into R and processed using limma and edgeR. Genes with minimal expression were removed, and normalization was performed using the TMM method followed by voom transformation. Differential expression analysis between *KRAS*-mutant and wild-type groups was performed using limma’s linear modeling and empirical Bayes shrinkage. Functional enrichment was conducted using clusterProfiler, including Gene Ontology (GO) analysis.

### Cell Culture

Mutant *KRAS* PDAC cell lines PANC-1 and MIAPaCa-2 were cultured in DMEM with high glucose (Gibco, 11965092), 10% FBS, and 1x antibiotic/antimitotic (30-004-CI, Corning). BxPC3 were cultured in RPMI with L-glutamine (10-040-CV, Corning), 10% FBS, and 1x antibiotic/antimitotic. Cells were treated with inhibitors MRTX-1133 (100 nM, 36413, Cayman Chemical), AMG-510 (1000 nM, 29465, Cayman Chemical), Trametinib (100 nM, 16292, Cayman Chemical), or Wortmannin (100 nM, 10010591, Cayman Chemical) for 24 and 48 h. For RMC-7977 treatment, PANC-1 cells were plated in normal DMEM media conditions and allowed to adhere overnight. Cells were then treated daily with RMC-7977 (100 nmol, HY-156498, Med Chem Express), the tool compound of daraxonrasib, for 48 h. The media was then changed to Opti-MEM, reduced serum media (31985070, Gibco) while being treated again with RMC-7977 daily for 48 h prior to collection for flow cytometry. For quantitative real-time polymerase chain reaction (qRT-PCR), RNA was isolated using Quick-RNA MiniPrep kits (R1055, Zymo Research), and concentration was measured using NanoDrop 2000. qRT-PCR was performed using a Luna Universal One-Step RT-qPCR Kit (E3005, New England Laboratories) and analyzed using Bio-Rad CFX96 system and CFX Maestro 2.3. The primer sequences used are provided in **Supplementary Table 1**.

Murine cell lines (iKras#1 and iKras#2) were cultured in RPMI with L-glutamine, 10% FBS and 1x antibiotic/antimitotic with 1 µg/ml doxycycline (D43020, RPI Research Products International) for continuous *Kras^G12D^*induction. To limit oncogenic *Kras^G12D^*expression, doxycycline was removed for 48 h. After 48 h, RNA was extracted and qRT-PCR was performed.

Isogenic PANC-1 G12D and G12R cell lines were generated by Synthego for the McFall laboratory as described in previous studies^7,36^. Cells were cultured in DMEM supplemented with 10% FBS, penicillin (100 U/mL), streptomycin (100 μg/mL), and L-glutamine (2 mM) at 37°C with 5% CO_2_. All cell lines were mycoplasma-free (MycoAlert Detection Kit) and used within 11 passages of acquisition to maintain cellular integrity and validated with short tandem repeat (STR) profiling to authenticate cell line identity prior to distribution; isogenic *KRAS*-mutant derivatives were generated from this authenticated parental PANC-1 line by CRISPR-mediated knock-in (Synthego) and were verified by sanger sequencing.

### Western Blotting

PANC-1 Isogenic cells were treated with increasing concentrations of daraxonrasib, RMC-6236 (HY-148439, MedChemExpress), at concentrations indicated for 48 h. PANC-1, MIAPaCa-2 and BxPC3 cells were treated with MRTX-1133, AMG-510, or wortmannin as described above. Cell lysates were generated using lysis buffer (1862301, Thermo Fisher Scientific or 20-188, Millipore) containing a protease inhibitor cocktail (Cell Signaling Technology) and 1X EDTA (78429, Pierce) and incubated on ice for 1 h with intermittent vortexing. The total protein concentration was determined using the Pierce protein assay (Thermo Fisher Scientific) or Qubit Protein BR assay (A50669, Thermo) per manufacturer’s instructions. Protein samples (20–30 µg per lane) were prepared using LI-COR 4X Protein Sample Loading Buffer (928-40004, Li-Cor) supplemented with β-mercaptoethanol (1:10, v/v, β-mercaptoethanol:sample buffer) and heated at 95°C for 5 min. Protein samples were resolved by electrophoresis on either 12% SDS–polyacrylamide gels and electrophoretically transferred to polyvinylidene difluoride (PVDF) membranes (IPFL00010, Millipore) for 30 min at 25 V using the Trans-Blot Turbo Transfer System (RRID:SCR_023156, BioRad) or 10% polyacrylamide gel and transferred to a PVDF membrane for 90 min at 80V on ice. Blots were then blocked in Licor Blocking buffer (927-60001, Li-Cor) and probed with the indicated primary antibodies incubated at 4°C overnight. Fluorescent secondaries were incubated for 1 h at room temperature in Licor Blocking Buffer containing 0.02% SDS and 0.2% Tween-20. Bands were then visualized using the Li-Cor Odyssey M imaging station (RRID:SCR_025709, Li-Cor Biosystems). Details of all antibodies used are provided in **Supplementary Table 2**.

### RNA Sequencing

Isogenic PANC-1 cell lines bearing either a *KRAS* G12D or G12R mutation were treated for 48 h with RMC-6236 (HY-148439, MedChemExpress) at 100 nM before collection in TRIzol. A quality control analysis was performed on the RNA samples using the Agilent TapeStation to assess quality and an RNA Qubit assay was used to determine quantity. Poly(A) RNA enrichment was conducted using NEBNext Poly(A) mRNA Magnetic Isolation Module (NEB), and the sequencing library was constructed by using the NEBNext Ultra II RNA Library Prep Kit (NEB) (P/N: E7765L, NEB, Ipswich, MA) following the manufacturer’s instructions (250 ng input material). End repair, A-tailing, and adapter ligation were performed to generate the final cDNA library. The library quality was assessed using the TapeStation and quantified using a qPCR-based method with the KAPA Library Quantification Kit (P/N: KK4873) and the QuantStudio 12K instrument. Prepared libraries were pooled in equimolar ratios, and the resulting pool was subjected to cluster generation using the NovaSeq XP System, following the manufacturer’s protocols. 150 bp paired-end sequencing was performed on the NovaSeq XP platform targeting 50M reads per sample. Raw sequencing data (FASTQ files) obtained from NovaSeq XP were subjected to quality control analysis, including read quality assessment. Real Time Analysis Software (RTA) and NovaSeq Control Software (NCS) (1.8.0; Illumina) were used for base calling. MultiQC (v1.7; Illumina) was used for data quality assessments. Raw sequencing data (FASTQ files) were assessed for quality using FastQC. Trim Galore paired-end mode was used to trim adapter sequences and low-quality reads. Reads were aligned to the human reference genome (hg38) using Spliced Transcripts Alignment to a Reference (STAR). Following alignment, gene-level quantification was performed using featureCounts. GENCODE primary assembly annotation (v44) was used for read counting. Data analyses were executed using R (version 4.4.2, 2024-10-31 ucrt), tailored for a 64-bit Windows platform. Packages used in the study consisted of “DESeq2(1.46.0)”, “edgeR(4.4.2)”, “clusterProfiler(4.14.6)”, “limma(3.62.2)”. Raw count matrices were imported from processed FASTQ files for analysis using DESeq2. Before analysis, lowly expressed genes were filtered out using the filterByExpr function. A Wald test was used for statistical significance between differentially expressed genes, with p-values adjusted for multiple comparisons via Benjamini-Hochberg false discovery rate (FDR). For principal component analysis and downstream visualization, raw counts were normalized by applying a Variance Stabilizing Transformation (VST). Supervised heatmaps, featuring hierarchical clustering of both samples and genes, were generated using the pheatmap R package. Gene Ontology analysis was conducted on differentially expressed gene sets (FDR < 0.05 and log2|FC| > 1) using the clusterProfiler package.

### Flow Cytometry

After inhibitor treatments, cells were detached using TrypLE (12604013, Gibco), washed 1–2 times with PBS, and blocked on ice in 1% (w/v) BSA in PBS for 30 min. Cells were pelleted at 1,000 × g for 5 min at 4°C and incubated with human anti-CD73 diluted in 1% BSA/PBS for 30 min on ice. Following antibody incubation, cells were pelleted (1,000 × g, 5 min, 4°C), washed 2–3 times with PBS, and fixed in 4% paraformaldehyde (15700, Electron Microscopy Sciences) on ice for 15 min. Cells were then washed 2–3 times in PBS, resuspended in 1% BSA/PBS, and analyzed on a BD LSRFortessa flow cytometer (BD Biosciences).

### Genetic Mouse Models

Mice were housed in accordance with National Institutes of Health guidelines in American Association for Accreditation of Laboratory Animal Care-accredited facilities at either the Salk Institute, Vanderbilt University, or the University of Nebraska Medical Center. The Salk Institute, Vanderbilt University, or UNMC Institutional Animal Care and Use Committees (IACUC) approved all animal studies.

LSL-*Kras^G12D/+^*, *Krt19^CreERT^*, *Hnf1*β*-CreER^T2^* and *LSL-Kras^G12D/+^;Pdx1-Cre* mice used in this study have been previously described^37–40^. For the *Kras^G12D^;Hnf1*β*-CreER^T2^* and *Kras^G12D^;Krt19CreERT* models, adult mice were treated with caerulein by intraperitoneal injection at 250 µg/kg, twice daily, 5 days a week, for 4 weeks at which time they were either sacrificed for pancreas collection or received 5 mg of tamoxifen daily for 5 days for induction of *Kras^G12D^*expression with 1 month recovery prior to sacrifice and tissue collection.

### KPC Syngeneic Subcutaneous Tumor Model

For the co-targeting study, a total of 5 × 10^4^ *Pdx1-Cre;LSL-Kras^G12D/+^;LSL-Trp53^R^*^172^*^H/+^* (KPC) cells were suspended in a 1:1 mixture of PBS and Matrigel and injected subcutaneously into the flank of 8-week-old male C57BL/6 mice. The KPC cell line was originally generated in the Tuveson laboratory from pancreatic tumors arising in KPC mice, which spontaneously develop PDAC. Since the KPC cell line was derived from a male mouse, only male recipient mice were used in this study. Tumor volumes were monitored twice weekly, and tumor volume was calculated as (length × width²)/2 and expressed in mm³. Mice were euthanized on day 28 following tumor implantation, and tumors were harvested for downstream analyses.

### Drug Treatment

In the co-targeting study, Quemliclustat (HY-125286, MedChemExpress), daraxonrasib - RMC-6236 (HY-148439, MedChemExpress), and trametinib (HY-10999, MedChemExpress) were formulated in 10% dimethyl sulfoxide (DMSO; HY-Y0320, MedChemExpress) and 90% corn oil (C8267, Sigma-Aldrich). Treatment started 8 days after tumor implantation and continued once daily by oral gavage until euthanasia on day 28. Quemliclustat (n = 9), daraxonrasib (n = 7), and trametinib (n = 10) were administered at doses of 10, 20, and 1 mg/kg, respectively. For combination treatment groups, quemliclustat was combined with either daraxonrasib (n = 9) or trametinib (n = 9) in the same dosing formulation and administered concurrently by oral gavage at the same doses used for the corresponding single agent treatments. Vehicle treated mice (n = 10) received 10% DMSO and 90% corn oil according to the same dosing schedule.

### Histological Analyses

Murine tissues were cut into 5 μm sections, mounted, and stained by H&E, PAS/AB, or for IHC as previously described^10,17^. Briefly, sections were deparaffinized, rehydrated through a graded ethanol series, permeabilized, and subjected to heat mediated antigen retrieval. Following blocking of endogenous peroxidase activity and nonspecific binding, sections were incubated with primary antibodies overnight at 4°C, followed by the appropriate secondary antibodies. Signal was detected using an avidin-biotin complex and 3,3′-diaminobenzidine (DAB) substrate (SK-4100, Vector Laboratories), and sections were counterstained with hematoxylin, mounted, and visualized by light microscopy. PAS/AB staining was performed per the manufacturer’s instructions. Details of all antibodies used are provided in **Supplementary Table 2**.

### Immunohistochemistry Quantitative Image Analysis

IHC staining was quantified using ImageJ software (http://imagej.nih.gov/ij/) software. Stain separation was performed using the Colour Deconvolution2 plugin, and DAB-positive staining was quantified as the percentage of positively stained area per field. Approximately 10 representative fields were analyzed per tissue section (n = 3 mice per group for analysis). Statistical analyses were performed using GraphPad Prism 10 (GraphPad, San Diego, CA, USA). Comparisons between groups were conducted using the Mann–Whitney U test (two unpaired groups). Statistical significance was defined as p < 0.05.

## Data Availability Statement

The human cell line RNA sequencing and mutation data were downloaded from cBioPortal (ccle_broad_2019)^13–16^. The data generated in this study are available within the article and its supplementary data files or upon request from the corresponding authors.

## REFERENCES

1. Varghese AM, Perry MA, Chou JF, et al. Clinicogenomic landscape of pancreatic adenocarcinoma identifies KRAS mutant dosage as prognostic of overall survival. Nat Med. Feb 2025;31(2):466–477. doi:10.1038/s41591-024-03362-3

2. Wasko UN, Jiang J, Dalton TC, et al. Tumour-selective activity of RAS-GTP inhibition in pancreatic cancer. Nature. May 2024;629(8013):927–936. doi:10.1038/s41586-024-07379-z

3. Holderfield M, Lee BJ, Jiang J, et al. Concurrent inhibition of oncogenic and wild-type RAS-GTP for cancer therapy. Nature. May 2024;629(8013):919–926. doi:10.1038/s41586-024-07205-6

4. Wolpin BM, Park W, Garrido-Laguna I, et al. Daraxonrasib in Previously Treated Advanced RAS-Mutated Pancreatic Cancer. N Engl J Med. May 7 2026;394(18):1790–1802. doi:10.1056/NEJMoa2505783

5. O’Reilly EM, Wainberg ZA, Hendifar AE, et al. Daraxonrasib or Chemotherapy in Previously Treated Metastatic Pancreatic Cancer. N Engl J Med. Jul 23 2026;395(4):325–337. doi:10.1056/NEJMoa2605555

6. Aronchik I, Kar S, Zhuang Y, et al. Acquired resistance to the RAS(ON) multi-selective inhibitor daraxonrasib guides rational combination therapy strategies in pancreatic cancer. Nat Med. Aug 2026;32(8):2865–2877. doi:10.1038/s41591-026-04537-w

7. Dorbin D, Herrera J, Davidson R, et al. Allele-Specific Mechanisms Guide Salvage Therapy to Overcome Daraxonrasib Resistance in Pancreatic Cancer. Cancer Res. Aug 27 2026;doi:10.1158/0008-5472.CAN-26-1181

8. Orlen M, Vostrejs WP, Sor R, et al. T-cell Dependency of Tumor Regressions and Complete Responses with RAS(ON) Multi-selective Inhibition in Preclinical Models of Pancreatic Ductal Adenocarcinoma. Cancer Discov. Aug 4 2025;15(8):1697–1716. doi:10.1158/2159-8290.CD-24-1475

9. Allard B, Allard D, Buisseret L, Stagg J. The adenosine pathway in immuno-oncology. Nat Rev Clin Oncol. Oct 2020;17(10):611–629. doi:10.1038/s41571-020-0382-2

10. Faraoni EY, Singh K, Chandra V, et al. CD73-Dependent Adenosine Signaling through Adora2b Drives Immunosuppression in Ductal Pancreatic Cancer. Cancer Res. Apr 4 2023;83(7):1111–1127. doi:10.1158/0008-5472.CAN-22-2553

11. DelGiorno K. CD73 Is a Critical Immune Checkpoint in a Molecular Subtype of Pancreatic Cancer. Cancer Res. Apr 4 2023;83(7):977–978. doi:10.1158/0008-5472.CAN-23-0268

12. Wainberg ZA, Manji GA, Bahary N, et al. Quemliclustat and chemotherapy with or without zimberelimab in metastatic pancreatic adenocarcinoma: a randomized phase 1 trial. Nat Med. Apr 2026;32(4):1267–1277. doi:10.1038/s41591-026-04283-z

13. Ghandi M, Huang FW, Jane-Valbuena J, et al. Next-generation characterization of the Cancer Cell Line Encyclopedia. Nature. May 2019;569(7757):503–508. doi:10.1038/s41586-019-1186-3

14. de Bruijn I, Kundra R, Mastrogiacomo B, et al. Analysis and Visualization of Longitudinal Genomic and Clinical Data from the AACR Project GENIE Biopharma Collaborative in cBioPortal. Cancer Res. Dec 1 2023;83(23):3861–3867. doi:10.1158/0008-5472.CAN-23-0816

15. Gao J, Aksoy BA, Dogrusoz U, et al. Integrative analysis of complex cancer genomics and clinical profiles using the cBioPortal. Sci Signal. Apr 2 2013;6(269):pl1. doi:10.1126/scisignal.2004088

16. Cerami E, Gao J, Dogrusoz U, et al. The cBio cancer genomics portal: an open platform for exploring multidimensional cancer genomics data. Cancer Discov. May 2012;2(5):401–4. doi:10.1158/2159-8290.CD-12-0095

17. Ma Z, Lytle NK, Chen B, et al. Single-Cell Transcriptomics Reveals a Conserved Metaplasia Program in Pancreatic Injury. Gastroenterology. Feb 2022;162(2):604–620 e20. doi:10.1053/j.gastro.2021.10.027

18. Takaori A, Hashimoto D, Ikeura T, et al. Impact of neoadjuvant therapy on gut microbiome in patients with resectable/borderline resectable pancreatic ductal adenocarcinoma. Pancreatology. Jun 2023;23(4):367–376. doi:10.1016/j.pan.2023.04.001

19. Minaei E, Joyce P, Wade SJ, et al. Gut microbial diversity is preserved through localised chemo-immunotherapy delivery in a KPC mouse model of pancreatic cancer. J Control Release. Oct 10 2025;386:114143. doi:10.1016/j.jconrel.2025.114143

20. Tintelnot J, Xu Y, Lesker TR, et al. Microbiota-derived 3-IAA influences chemotherapy efficacy in pancreatic cancer. Nature. Mar 2023;615(7950):168–174. doi:10.1038/s41586-023-05728-y

21. Mirji G, Worth A, Bhat SA, et al. The microbiome-derived metabolite TMAO drives immune activation and boosts responses to immune checkpoint blockade in pancreatic cancer. Sci Immunol. Sep 9 2022;7(75):eabn0704. doi:10.1126/sciimmunol.abn0704

22. Kao DJ, Saeedi BJ, Kitzenberg D, et al. Intestinal Epithelial Ecto-5’-Nucleotidase (CD73) Regulates Intestinal Colonization and Infection by Nontyphoidal Salmonella. Infect Immun. Oct 2017;85(10)doi:10.1128/IAI.01022-16

23. Zhang J, Dong H, Liang L, et al. Targeting gut microbiota and arginase boosts MEK inhibitors’ enhancement of antitumour immunity via MHC-I upregulation in colorectal cancer. Br J Cancer. Oct 2025;133(6):809–822. doi:10.1038/s41416-025-03106-1

24. Tang T, Huang X, Lu M, Zhang G, Han X, Liang T. Transcriptional control of pancreatic cancer immunosuppression by metabolic enzyme CD73 in a tumor-autonomous and -autocrine manner. Nat Commun. Jun 8 2023;14(1):3364. doi:10.1038/s41467-023-38578-3

25. Li Y, He Y, Peng J, et al. Mutant Kras co-opts a proto-oncogenic enhancer network in inflammation-induced metaplastic progenitor cells to initiate pancreatic cancer. Nat Cancer. Jan 2021;2(1):49–65. doi:10.1038/s43018-020-00134-z

26. Li AL, Sugiura K, Nishiwaki N, et al. FRA1 controls acinar cell plasticity during murine Kras(G12D)-induced pancreatic acinar to ductal metaplasia. Dev Cell. Nov 18 2024;59(22):3025–3042 e7. doi:10.1016/j.devcel.2024.07.021

27. Reinhardt J, Landsberg J, Schmid-Burgk JL, et al. MAPK Signaling and Inflammation Link Melanoma Phenotype Switching to Induction of CD73 during Immunotherapy. Cancer Res. Sep 1 2017;77(17):4697–4709. doi:10.1158/0008-5472.CAN-17-0395

28. Arnold F, Del Vecchio A, Hussain Z, Sherman MH. Heterocellular crosstalk and architecture of the pancreatic tumour microenvironment. Nat Rev Cancer. Apr 2026;26(4):285–304. doi:10.1038/s41568-025-00905-9

29. Mahadevan KK, Maldonado AS, Li B, et al. Oncogenic Kras targeting with MRTX1133 or Daraxonrasib specifically synergize with anti-CTLA4 to promote anti-tumor immunity in pancreatic cancer. Nat Commun. Jul 27 2026;17(1)doi:10.1038/s41467-026-75960-3

30. McAndrews KM, Mahadevan KK, Li B, et al. CDK8 remodels the tumor microenvironment and promotes resistance to KRAS(G12D) inhibitors and daraxonrasib in PDAC. EMBO J. Aug 2026;45(16):5727–5758. doi:10.1038/s44318-026-00854-5

31. Kemp SB, Cheng N, Markosyan N, et al. Efficacy of a Small-Molecule Inhibitor of KrasG12D in Immunocompetent Models of Pancreatic Cancer. Cancer Discov. Feb 6 2023;13(2):298–311. doi:10.1158/2159-8290.CD-22-1066

32. Velez-Delgado A, Donahue KL, Brown KL, et al. Extrinsic KRAS Signaling Shapes the Pancreatic Microenvironment Through Fibroblast Reprogramming. Cell Mol Gastroenterol Hepatol. 2022;13(6):1673–1699. doi:10.1016/j.jcmgh.2022.02.016

33. Liang Y, Du M, Li X, et al. Upregulation of Lactobacillus spp. in gut microbiota as a novel mechanism for environmental eustress-induced anti-pancreatic cancer effects. Gut Microbes. Dec 2025;17(1):2470372. doi:10.1080/19490976.2025.2470372

34. Han ZY, Fu ZJ, Wang YZ, et al. Probiotics functionalized with a gallium-polyphenol network modulate the intratumor microbiota and promote anti-tumor immune responses in pancreatic cancer. Nat Commun. Aug 17 2024;15(1):7096. doi:10.1038/s41467-024-51534-z

35. Mager LF, Burkhard R, Pett N, et al. Microbiome-derived inosine modulates response to checkpoint inhibitor immunotherapy. Science. Sep 18 2020;369(6510):1481–1489. doi:10.1126/science.abc3421

36. Kamgar M, Hobbs GA, Davidson R, et al. Mutant and Wild-Type RAS Cross-talk and Stoichiometric Deficiencies Are Determinants of Sensitivity to Targeted Therapies in KRASG12R Pancreatic Ductal Adenocarcinoma. Cancer Res. Apr 15 2026;86(8):2042–2059. doi:10.1158/0008-5472.CAN-25-0018

37. Jackson EL, Willis N, Mercer K, et al. Analysis of lung tumor initiation and progression using conditional expression of oncogenic K-ras. Genes Dev. Dec 15 2001;15(24):3243–8. doi:10.1101/gad.943001

38. Means AL, Xu Y, Zhao A, Ray KC, Gu G. A CK19(CreERT) knockin mouse line allows for conditional DNA recombination in epithelial cells in multiple endodermal organs. Genesis. Jun 2008;46(6):318–23. doi:10.1002/dvg.20397

39. Solar M, Cardalda C, Houbracken I, et al. Pancreatic exocrine duct cells give rise to insulin-producing beta cells during embryogenesis but not after birth. Dev Cell. Dec 2009;17(6):849–60. doi:10.1016/j.devcel.2009.11.003

40. Strickland LN, Liu W, Hussein U, et al. Preventive Treatment with a CD73 Small Molecule Inhibitor Enhances Immune Surveillance in K-Ras Mutant Pancreatic Intraepithelial Neoplasia. Cancer Prev Res (Phila). Oct 1 2024;17(10):457–470. doi:10.1158/1940-6207.CAPR-24-0200

