## Supplemental Figures for "Oncogenic KRAS-MAPK signaling sustains a CD73 dependent therapeutic vulnerability in pancreatic cancer"

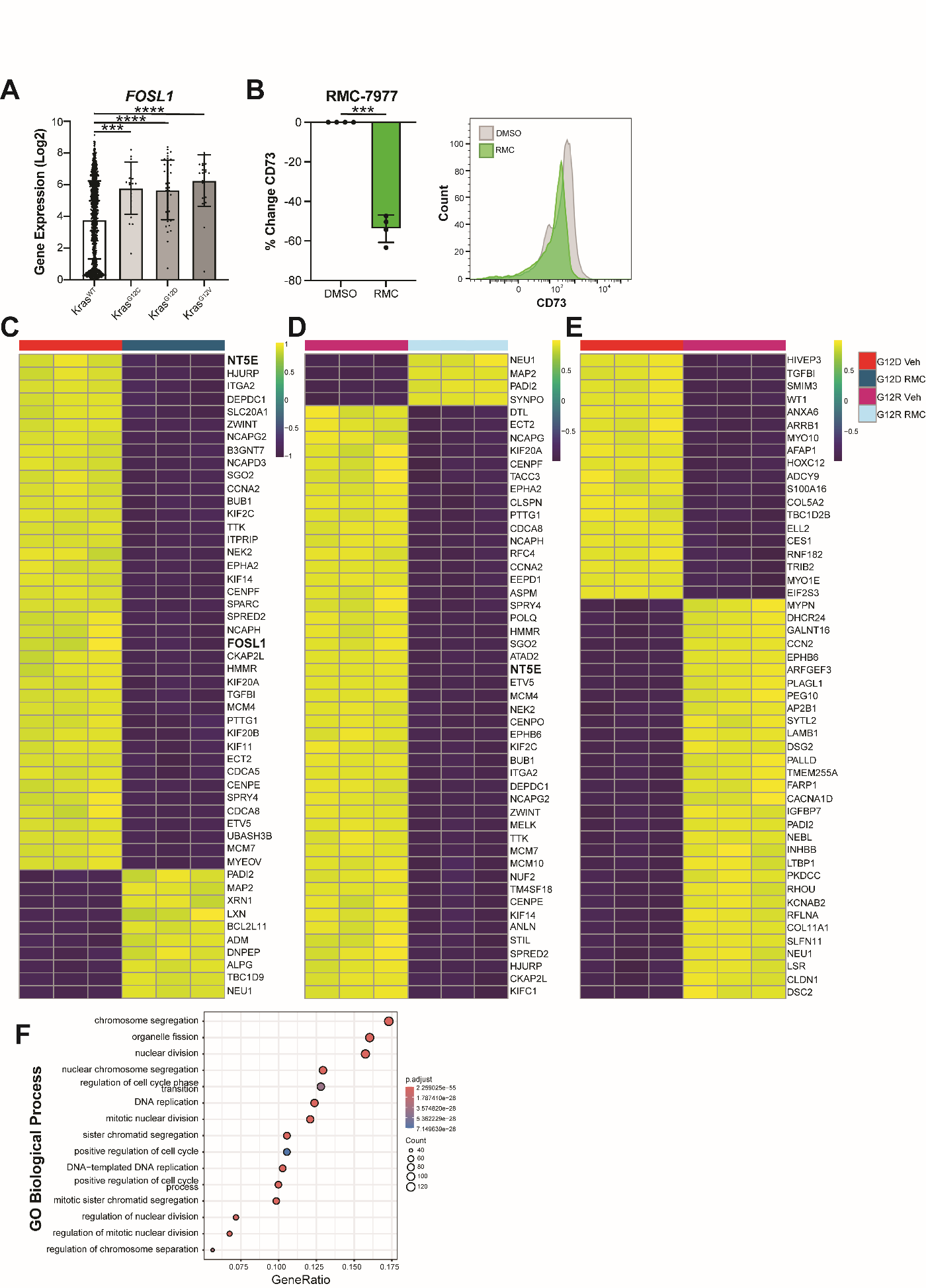
**SUPPLEMENTARY FIGURES**

**Supplementary Figure 1. CD73 expression is driven by oncogenic KRAS.** (**A**) Expression of known KRAS target gene *FOSL1* in cBioPortal cell line RNA-seq data stratified by *KRAS* mutation status. (**B**) CD73 expression in the PANC-1 (G12D) cell line treated with control or daraxonrasib tool compound RMC-7977, measured by flow cytometry. Heatmaps of gene expression comparing control vs daraxonrasib treated (**C**) PANC-1 G12D or (**D**) PANC-1 G12R cell lines or directly comparing (**E**) the control-treated PANC-1 G12D and G12R cell lines. (**F**) GO analysis showing a loss of proliferation-associated pathway gene expression in PANC-1 G12R cells with daraxonrasib treatment. Statistical significance was determined by a Welch’s $t$-test for two-group comparisons or one-way ANOVA followed by Dunnett’s post-hoc test for multiple comparisons. ***p < 0.001; ****p < 0.0001.

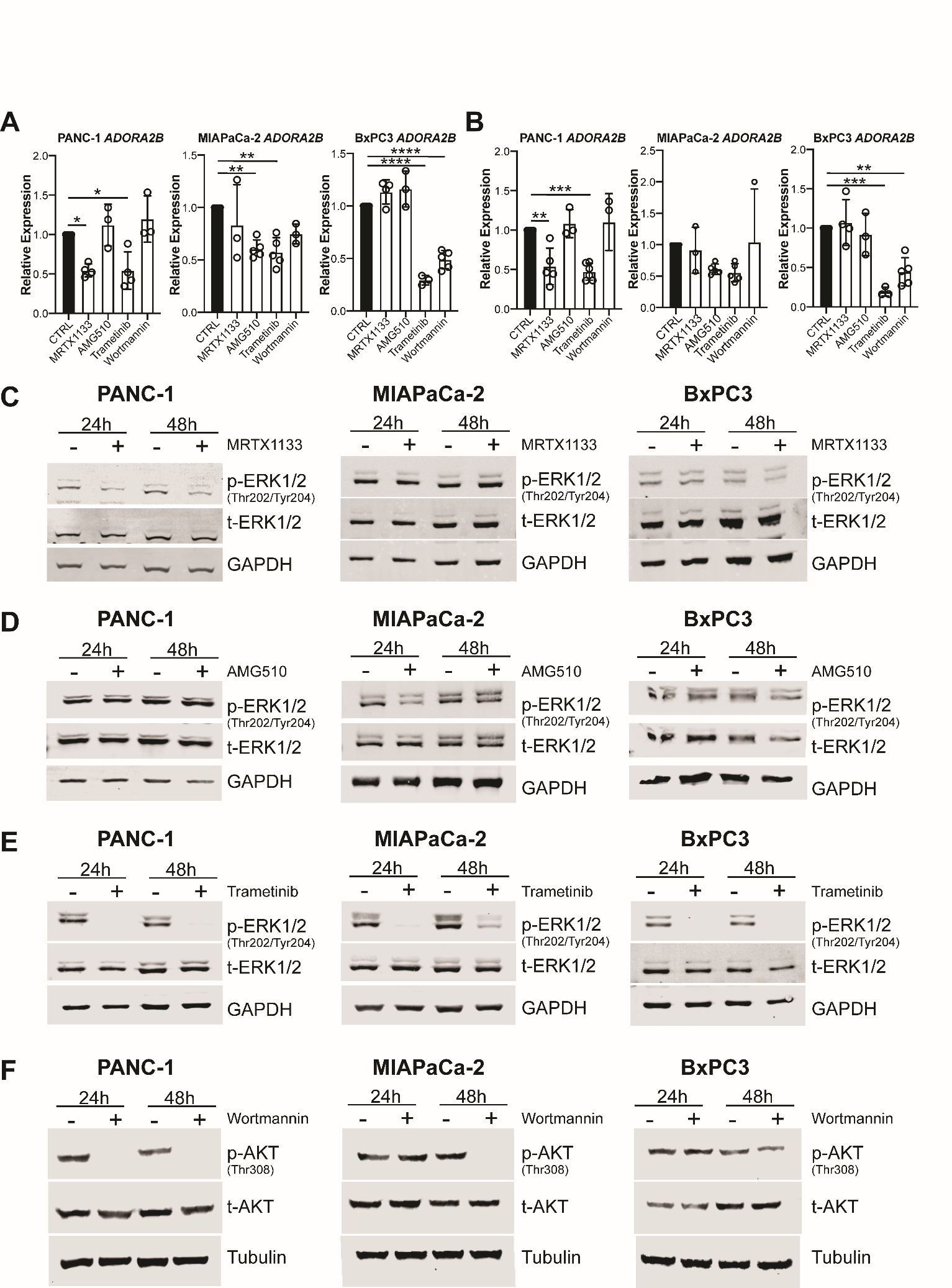
**Supplementary Figure 2. Oncogenic KRAS drives expression of adenosine receptor *ADORA2B* in pancreatic cancer cell lines.** Expression of *ADORA2B* in PANC-1 (*KRAS^G12D^*), MIAPaCa-2 (*KRAS^G12C^*), or BxPC3 (*KRAS* wild-type) pancreatic cancer cell lines treated with *KRAS^G12D^* inhibitor MRTX1133, *KRAS^G12C^* inhibitor AMG510, MEK inhibitor trametinib, or PI3K inhibitor wortmannin for either (**A**) 24 or (**B**) 48 h showing a loss of *ADORA2B* in PANC-1 or MIAPaCa-2 cell lines with KRAS or MEK inhibition and in BxPC3 lines with MEK or PI3K inhibition. (**C**) Western Blots showing a loss of pERK1/2 ^Thr202+Tyr204^ within PANC-1 cells, but not MIAPaCa-2 or BxPC3 with MRTX-1133 treatment, (**D**) a loss of pERK1/2 in MIAPaCa-2 cells, but not PANC-1 or BxPC3 with AMG510 treatment, (**E**) a loss of pERK1/2 in all cell lines with trametinib treatment, and (**F**) a loss of pAKT with wortmannin treatment. Statistical significance was determined by a one-way ANOVA followed by Dunnett’s post-hoc test. *p < 0.05; **p < 0.01; ***p < 0.005; ****p < 0.001.

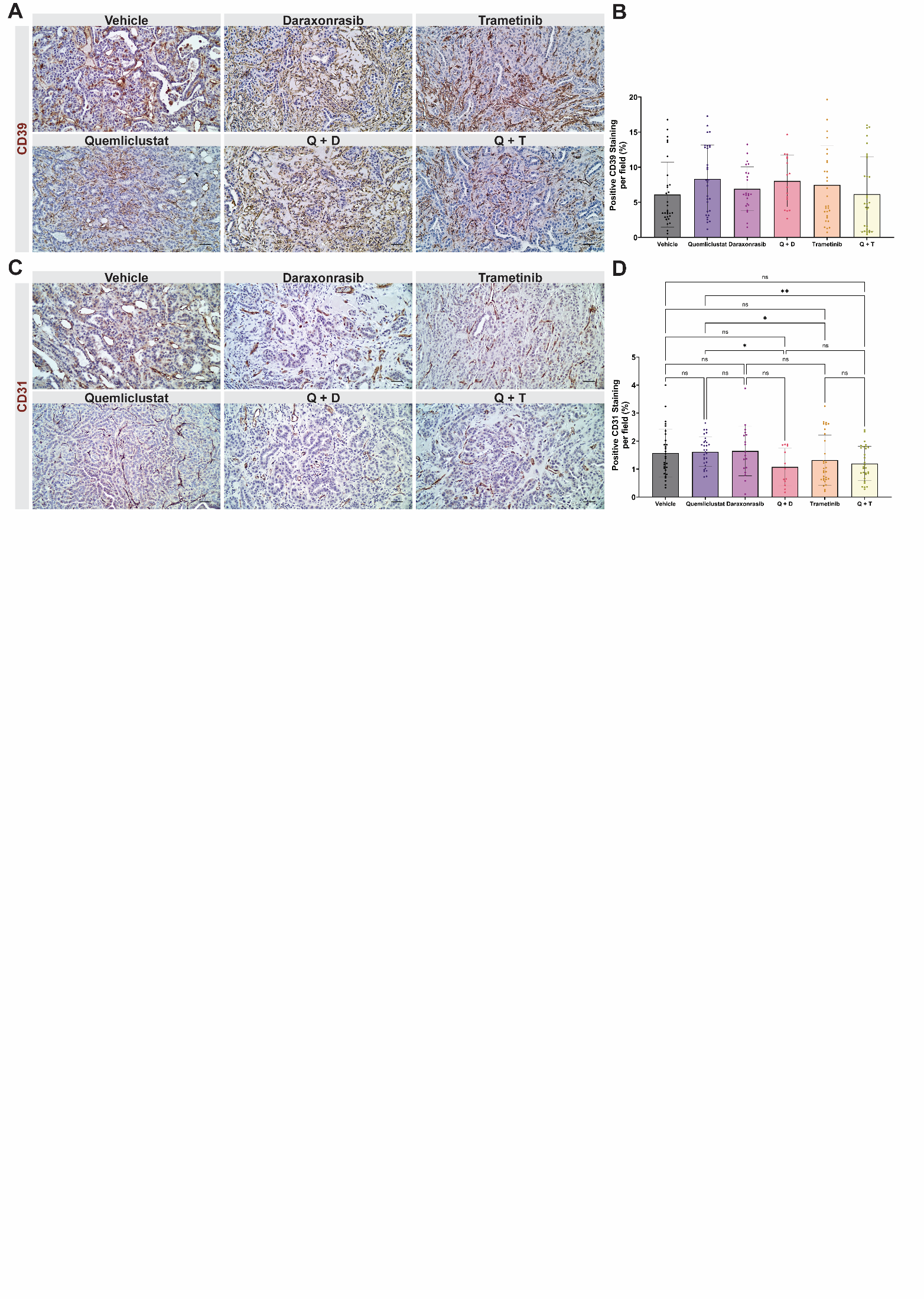
**Supplementary Figure 3. CD73 and KRAS-MAPK pathway inhibition alters tumor vascularization with limited effects on CD39 expression.** Representative images of (**A**) CD39 and (**C**) CD31 IHC staining of mouse subcutaneous tumor sections from the different treatment groups. Scale bars are 50 μm. Quantification of percent positive of (**B**) CD39 (ns between all groups) and (**D**) CD31 staining per field in each group. All quantifications were performed for n = 3 mice per treatment group, and 5–10 fields per mouse were analyzed. Statistical significance was determined by Mann–Whitney U test. ns, not significant; *p < 0.05; **p < 0.01.

| **Gene** | **Foward Primer** | **Reverse Primer** |
| --- | --- | --- |
| *RPLP0* | GCC AAT AAG GTG CCA GCT G | CTC CCA CCT TGT CTC CAG TC |
| *NT5E* | CCC ATT CTT CTA AAC AGC AGC ATT C | TGA GAG GAG CCA TCC AGA TAG AC |
| *FOSL1* | CGC GAG CGG AAC AAG CT | GTC TCC GCC TGC AGG AAG |
| *ADORA2B* | ACG GCT GCC TCT TCC TCG | GGA CCC AGA GGA CAG CAA TGA |
| *Nt5e* (mouse) | GCA AAC ATT AAG GCA CGG GG | TCA CCG CCA ACA GAG AGA AC |

**Supplementary Table 1. Primer sequences.**

| **Antibody** | **Dilution** | **Source** | **Identifier** |
| --- | --- | --- | --- |
| **IHC** | | | |
| CD73 | 1:100-200 | CST | 13160 |
| FRA1 | 1:50 | CST | 28801 |
| pERK1/2 (T202/T185) | 1:500 | abcam | ab214036 |
| Ki67 | 1:200 | abcam | ab15580 |
| αSMA | 1:500 | abcam | ab5694 |
| CD8α | 1:250 | abcam | ab217344 |
| Granzyme B | 1:200 | abcam | ab255598 |
| CD68 | 1:100 | abcam | ab283654 |
| CD163 | 1:500 | abcam | ab182422 |
| CD39 | 1:200 | abcam | ab223842 |
| CD31 | 1:500 | abcam | ab281583 |
| MUC5AC | 1:500 | Invitrogen | MA5-12178 |
| **Flow Cytometry** | | | |
| FITC anti-human CD73 | 1:100 | BioLegend | 344015 |
| **Western Blotting** | | | |
| pERK1/2 (Thr202/Tyr204) | 1:1000 | BioLegend | 675502 |
| ERK1/2 | 1:1000 | BioLegend | 686902 |
| alpha/beta-Tubulin | 1:1000 | CST | 2148 |
| CD73 | 1:1000 | CST | 13160 |
| p44/42 MAPK  (Erk1/2) (Thr202/Tyr204) | 1:1000 | CST | 9101 |
| p-p44/42 MAPK | 1:1000 | CST | 4696 |
| pAKT (Thr308) | 1:1000 | CST | 4056 |
| AKT (pan) | 1:1000 | CST | 2920 |
| Rabbit IR800 | 1:5000 | Li-cor | 925-32211 |
| Mouse IR680 | 1:5000 | Li-cor | 925-68070 |
| GAPDH 488 | 1:500 | Invitrogen | MA5-15738-A488 |

**Supplementary Table 2. Antibodies used.**

**Supplementary File 1: Differentially Expressed Genes between *KRAS* wild-type and mutant cancer cell lines in cBioPortal.**

**Supplementary File 2: Differentially Expressed Genes between control and daraxonrasib treated *KRAS^G12D^* PANC-1 cells.**

**Supplementary File 3: Differentially Expressed Genes between control and daraxonrasib treated *KRAS^G12R^* PANC-1 cells.**
